# MINERVA: A facile strategy for SARS-CoV-2 whole genome deep sequencing of clinical samples

**DOI:** 10.1101/2020.04.25.060947

**Authors:** Chen Chen, Jizhou Li, Lin Di, Qiuyu Jing, Pengcheng Du, Chuan Song, Jiarui Li, Qiong Li, Yunlong Cao, X. Sunney Xie, Angela R. Wu, Hui Zeng, Yanyi Huang, Jianbin Wang

## Abstract

The novel coronavirus disease 2019 (COVID-19) pandemic poses a serious public health risk. Analyzing the genome of severe acute respiratory syndrome coronavirus 2 (SARS-CoV-2) from clinical samples is crucial for the understanding of viral spread and viral evolution, as well as for vaccine development. Existing sample preparation methods for viral genome sequencing are demanding on user technique and time, and thus not ideal for time-sensitive clinical samples; these methods are also not optimized for high performance on viral genomes. We have developed <u>M</u>etagenom<u>I</u>c R<u>N</u>A <u>E</u>n<u>R</u>ichment <u>V</u>ir<u>A</u>l sequencing (MINERVA), a facile, practical, and robust approach for metagenomic and deep viral sequencing from clinical samples. This approach uses direct tagmentation of RNA/DNA hybrids using Tn5 transposase to greatly simplify the sequencing library construction process, while subsequent targeted enrichment can generate viral genomes with high sensitivity, coverage, and depth. We demonstrate the utility of MINERVA on pharyngeal, sputum and stool samples collected from COVID-19 patients, successfully obtaining both whole metatranscriptomes and complete high-depth high-coverage SARS-CoV-2 genomes from these clinical samples, with high yield and robustness. MINERVA is compatible with clinical nucleic extracts containing carrier RNA. With a shortened hands-on time from sample to virus-enriched sequencing-ready library, this rapid, versatile, and clinic-friendly approach will facilitate monitoring of viral genetic variations during outbreaks, both current and future.

## Introduction

As of May 22, 2020, the ongoing COVID-19 viral pandemic has affected 5 million people in over 200 countries and territories around the world, and has claimed more than 320 thousand lives^1^. Closely monitoring the genetic diversity and distribution of viral strains at the population level is essential for epidemiological tracking, and for understanding viral evolution and transmission; additionally examining the viral heterogeneity within a single individual is imperative for diagnosis and treatment^2^. The disease-causing pathogen, severe acute respiratory syndrome coronavirus 2 (SARS-CoV-2), was identified from early disease cases and its draft genome sequenced within weeks, thanks to the rapid responses from researchers around the world^3–6^. The initial SARS-CoV-2 draft genome was obtained independently from the same early COVID-19 patient samples using various conventional RNA-seq sequencing library construction methods. Although these library construction methods successfully generated a draft genome, several drawbacks hinder the use of these methods for routine viral genome sequencing from the surge of clinical samples during an outbreak.

One direct library construction approach which was used to generate the SARS-CoV-2 draft genome^3–6^ essentially captures each sample’s entire metatranscriptome, in which SARS-CoV-2 is just one species among many. The abundance of SARS-CoV-2 in clinical swabs, sputum, and stool samples is often low^2^,^7^, therefore this catch-all method requires deeper sequencing of each sample in order to obtain sufficient coverage and depth of the whole viral genome, which increases the time and cost of sequencing. Target enrichment with spiked-in primers can improve SARS-CoV-2 genome coverage^8^, but the reliance on specific primers inherently limits this approach for the profiling of fast evolving viruses such as coronaviruses. The same limitation applies to multiplex RT-PCR-based strategies^9^. Additionally, once the sample is subject to targeted amplification during the initial reverse transcription (RT) steps, its metatranscriptomic information is lost forever.

Currently, the most comprehensive strategy is the combination of metatranscriptomics profiling with post-library SARS-CoV-2 target enrichment^9^. However, in most conventional RNA-seq methods, the double-strand DNA ligation (dsDL) portion of the protocol is usually the most demanding on hands-on time and user technique^10^. When superimposed on the target enrichment process, these labor-intensive and lengthy protocols become impractical for routine use in the clinic, much less for the timely monitoring of viral genetics and evolution on large volumes of samples during an outbreak. Furthermore, due to the low molecular efficiency of dsDL, these protocols also require a high amount of input material, further restricting their application on clinical samples.

Summarily, although next generation sequencing platforms are high-throughput and have short turn-around time, library construction from samples – whether including targeted enrichment or not – remains a major bottleneck. To broadly apply viral sequencing on clinical samples, especially during outbreaks when biomedical resources are already limited, a rapid, simple, versatile, and scalable sample library construction method that does not compromise on performance is urgently needed.

Recently, we reported a new RNA-seq library construction strategy that aims to address some of these challenges: SHERRY avoids the problematic dsDL step in library construction by taking advantage of the newly discovered Tn5 tagmentation activity on RNA/DNA hybrids, to directly tag RNA/cDNA fragments with sequencing adapters^10^. As such, SHERRY has minimal sample transfers and greatly reduced hands-on time, making it simple, robust, and suitable for inputs ranging from single cells to 200 ng total RNA. We now combine the advantages of a tailored SHERRY protocol, which improved coverage of whole metatranscriptome, with a simplified post-library target enrichment protocol. <u>M</u>etagenom<u>I</u>c R<u>N</u>A <u>E</u>n<u>R</u>ichment <u>V</u>ir<u>A</u>l sequencing or MINERVA, is an easy-to-use, versatile, scalable, and cost-effective protocol that yields high-coverage high-depth SARS-CoV-2 genome, while preserving the sample’s rich metatranscriptomic profile. The hands-on time required from clinical sample to sequencing-ready library using conventional approaches without enrichment is 190 min; MINERVA requires only 100 min hands-on time, and if deep viral coverage is desired, an additional 90 min for post-library enrichment, totaling 190 min for the entire workflow (**Fig. S1**), making MINERVA practical for high-volume, routine clinical use. We applied MINERVA to various types of COVID-19 samples and successfully obtained up to 10,000-fold SARS-CoV-2 genome enrichment.

This strategy will facilitate all studies regarding SARS-CoV-2 genetic variations in the current pandemic, and can also be applied to other pathogens of interest.

## Results

### Metagenomic RNA enrichment viral sequencing (MINERVA)

To analyze both metagenomics and SARS-CoV-2 genetics from COVID-19 patient samples, we developed a two-stage metagenomic RNA enrichment viral sequencing strategy termed MINERVA (**Fig. 1A**). First, we employed a SHERRY-based RNA-seq pipeline for metagenomic analysis. Since clinical samples may contain DNA, RNA, and possibly carrier RNA, MINERVA starts with ribosomal RNA (rRNA) removal and optional simultaneous carrier RNA removal, followed by DNase I treatment. The remaining RNA is then subject to standard SHERRY. Previously we observed 3’ bias in SHERRY libraries; to address this, we used 10 ng mouse 3T3 cell total RNA as starting material, and tested whether adding random decamers (N10) during RT could improve coverage evenness (**Fig. S2**). Compared with the standard SHERRY protocol, which uses 1 μM T30VN primer during RT, the supplement of 1 μM N10 indeed improves gene body coverage evenness, presumably by improving the RT efficiency. When the N10 concentration was further increased to 10 μM, we observed almost no coverage bias in the gene body. The high N10 concentration can result in an increased rRNA ratio in the product, sometimes as high as 90%, but MINERVA employs rRNA removal as the first step prior to RT, thus negating this problem. We also performed enzyme titration with homemade and commercial Tn5 transposomes. Based on these N10 and Tn5 titration results, we used 10 μM N10 during RT and 0.5 μl V50 for each 20-μl tagmentation reaction in all following experiments. The whole procedure from nucleic acid to metagenomic sequencing-ready library, including wait time, takes 5.5 hours (**Fig. S1**).

**Figure 1.**
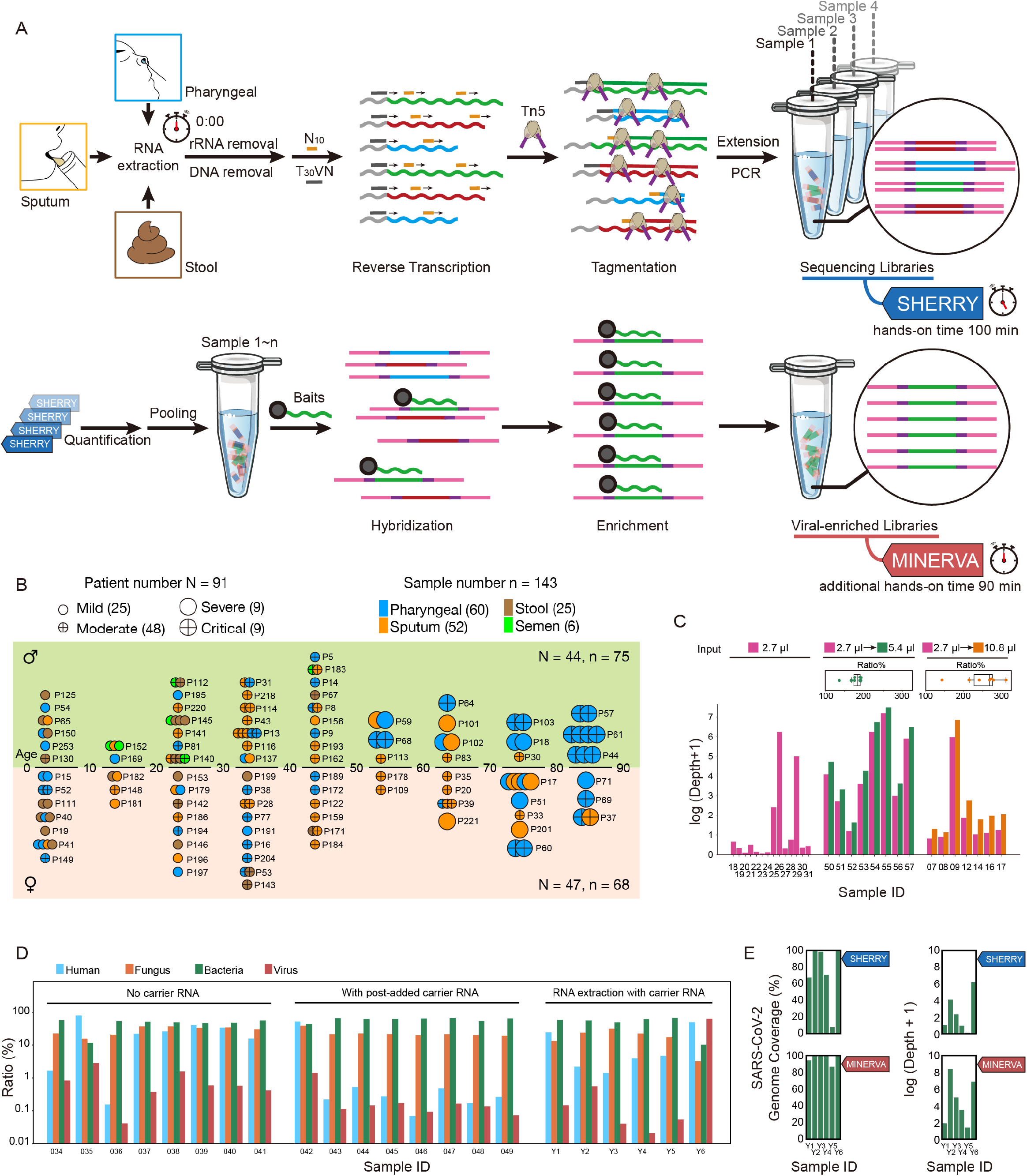
Scheme and optimization of MINERVA. (A) RNA extracted from pharyngeal swabs, sputum and stool samples undergo rRNA and DNA removal before a SHERRY processing pipeline metagenomic sequencing library construction. Multiple libraries were then pooled for SARS-CoV-2 sequence enrichment. (B) COVID-19 sample profiles, showing the age group, sex, severity, and re-sampling status of each patient. (C) Effect of sample input and reaction volume on sequencing depth of SARS-CoV-2 genome. (D) Metagenomic results of carrier RNA removal tests. (E) SARS-CoV-2 results of carrier RNA removal test.

For target enrichment, we first quantified SARS-CoV-2 abundance in each metagenomic sequencing library using an N gene qPCR assay, and pooled eight libraries based on quantification results. Then we performed standard in-solution hybridization on the pooled library with biotinylated RNA probes covering the whole viral genome. The enrichment procedure takes 7~13 hours; the entire MINERVA pipeline can be completed within 12~18 hours.

### MINERVA is compatible with COVID-19 samples

To evaluate its performance on clinical samples, we applied MINERVA on 143 samples collected from 91 COVID-19 patients, with samples types including pharyngeal swabs, sputum, stool, and semen. These patients were admitted to Ditan Hospital within a three-month period from January to April 2020, presenting different symptom severity (**Fig. 1B and Table S1-S3**). Some patients were re-sampled longitudinally to investigate temporal and intra-host viral heterogeneity. We first tested the effect of sample input volume on MINERVA results. Using just 2.7-ul of sample input led to satisfactory SARS-CoV-2 coverage, and scaling up the reaction volume to 5.4-ul further improved the MINERVA data quality (**Fig. 1C**). Using the same samples and at the same sequencing depth, more input in a higher reaction volume generated deeper SARS-CoV-2 genome coverage.

Carrier RNA, which is widely used in viral DNA/RNA extraction before RT-qPCR assays, severely impacts high-throughput sequencing analysis. Therefore, most RT-qPCR positive clinical samples are not amenable to further viral genetic studies. We explored the effect of adding polyT oligos during the rRNA removal step to simultaneously remove spike-in polyA RNA and carrier RNA. By incorporating this step in MINERVA, we successfully avoided the overwhelming representation of unwanted RNA sequences while retaining desired metagenomic and SARS-CoV-2 information (**Fig. 1D and 1E**).

### MINERVA captures metagenomic signatures of COVID-19 samples

We benchmarked MINERVA against conventional dsDL strategies in head-to-head comparisons of the first 79 clinical samples sequenced. On average, we sequenced 1-3 Gbp for each MINERVA library, and nearly 100 Gbp for each dsDL library (**Fig. S3**). The metagenomic compositions of SHERRY and dsDL libraries were comparable: total virus, fungus, and bacteria ratios were highly concordant between the two methods (**Fig. S4**); bacterial heterogeneity as measured by entropy is also correlated between the two.

We performed various analyses to explore the metagenomic composition of different samples types, and to assess whether metagenomic signatures correlate with disease severity. First and foremost, we observed that the metagenomic composition of different sample types show body site-specific features. Principle components analysis of bacterial sequences showed a clear separation between stool samples and the other sample types along PC1 (**Fig. 2A**), and this is reflected in analysis at both the genus and species levels, conveying the unique microbial environment of this body site. This phenomenon is most prominently reflected by the bacterial composition, but is also somewhat reflected in the viral composition (**Fig. S5**). We then identified the specific microbes that drive this separation of sample types, and found some microbes to be body site-specific. For example, stool samples contained *Bacteroides*, whereas the pharyngeal and sputum samples were rich in *Streptococcus* (**Fig. 2B and S5**); a few samples are highly abundant in known pathogenic species such as *Candida*, which is only found in orally obtained samples (**Fig. S5**). There also appears to be separation between samples by COVID-19 symptom severity along PC2 (**Fig. 2A**), which is supported by our analysis of specific microbial species (**Fig. 2B**). We found the bacterial metagenomic signature could be used to cluster most of the samples from “Critical” patients: samples from severe and critical condition patients are abundant in *Pseudomonas*, whereas *Streptococcus* is abundant in less severe condition samples.

**Figure 2.**
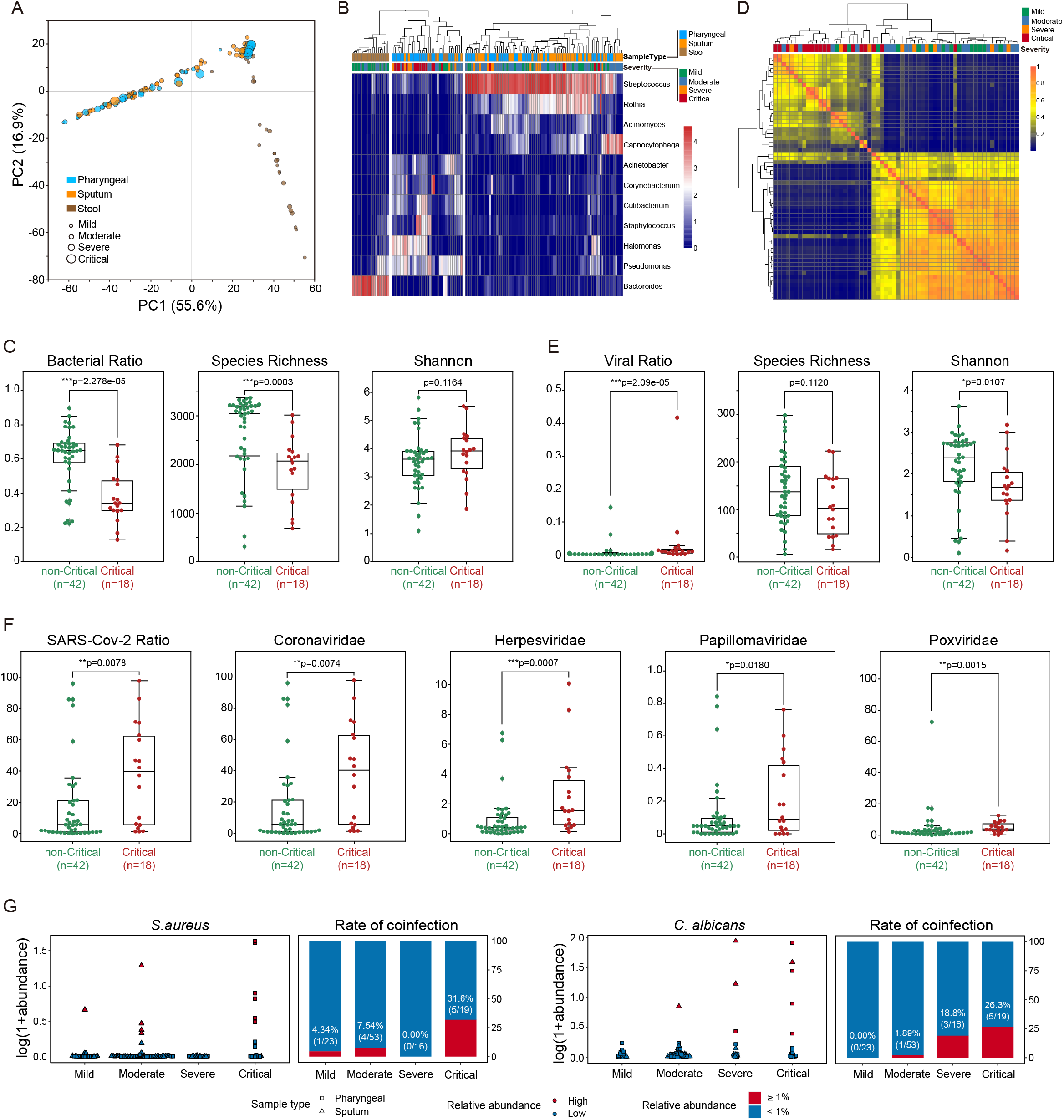
Metagenomic analyses of different sample types using MINERVA. (A) PCA analysis of bacterial composition in different sample types reveal body site-specific features. PCA analysis is based on bacterial genus of different sample types (60 pharyngeal, 51 sputum, and 25 stool samples). The bacterial genus composition of pharyngeal and sputum samples are more similar to each other, while stool samples are distinct from all other sample types. (B) Clustering analysis of bacterial composition in different sample types reveals characteristic microbial features in patients with the most severe disease symptoms. Bacteroides was dominant in stool samples while in oral samples (pharyngeal and sputum), samples from critical patients can be easily distinguished from patients with lower disease severity by the low abundance of Streptococcus and Rothia, and the high abundance of Pseudomonas. (C) Bacterial abundance and composition in pharyngeal swab samples significantly distinguish between critical and non-critical patients. Comparison of bacterial ratio and within-subject diversity between critical (n=42) and non-critical (n=18) pharyngeal samples. The bacterial ratio and species richness were significantly lower in Critical patients compared with non-Critical patients, while the Shannon index of alpha diversity is slightly higher, though not significant (Mann-Whitney U test). (D) Bacterial metagenomic composition is similar among critical patient samples and distinct from non-critical patient samples. Bray-Curtis beta diversity among all pharyngeal samples (n=60) at different stages. Severity samples were distinct from other stages. (E) Viral abundance correlates with disease severity. Comparison of viral ratio and within-subject diversity between critical samples (n=42) and non-critical pharyngeal samples (n=18). The viral ratio was significant higher in critical samples, while the species richness and Shannon index was slightly lower (Mann-Whitney U test). (F) Certain viral families correlate strongly with disease severity. Comparison of SARS-Cov-2 and other viral family relative abundance between critical (n=42) and non-critical (n=18) pharyngeal samples. The relative abundance of SARS-Cov-2, Coronaviridae, Herpesviridae, Papillomaviridae and Poxviridae were significantly higher in Critical samples (Mann-Whitney U test). (G) Metagenomic analysis of SHERRY libraries detect potential co-infection in specific patient samples. The cutoff of relative abundance for infection detection was set to 1% to avoid potential false positives. Higher rate of Staphylococcus aureus and Candida albicans can be detected in critical patients.

To further explore how bacterial composition reflects disease severity, we computed the bacterial ratio, bacterial species richness, and the Shannon Diversity Index for each sample type, and segmented samples by symptom severity (**Fig. S6**). Indeed, the bacterial abundance and composition in different sample types generally reflects disease severity. In particular pharyngeal swab samples show a statistically significant difference in bacterial ratio and species richness when comparing critical patients (“critical” group) with non-critical patients (“mild”, “moderate”, and “severe” groups) (**Fig. 2C**). The Shannon Diversity Index, however, is similar for all disease severities, indicating that although the overall bacterial abundance is reduced in critical patients, the relative abundance of different species remains stable. Interestingly, this phenomenon appears to also correlated with patient age – In elderly patients above the age of 60, both bacterial ratio and species richness are significantly reduced in critical patients as compared to non-critical; this trend is not observed in patients younger than 60 years of age (**Fig S6**). To further assess the relationship between bacterial metagenomic composition and disease severity, we calculated the pairwise Bray-Curtis similarity for pharyngeal swab samples, and found that mild, moderate, and severe patient samples are clustered together and intermixed, while critical samples cluster separately with each other (**Fig. 2D**). The bacterial metagenomic composition is similar among critical patients and suggests that they share common features distinct from non-critical patients. In light of recent studies of the role of host immunity responses in critically ill COVID-19 patients^11–14^, our observations of the metagenomic signature could be indicative of the systemic impact of the host immune response on commensal microbes, fungi, and other viruses in the body. Currently, the number of critical patient samples in the younger group is limited, but the trend is worth further investigation as more samples are collected over time.

In addition to the bacterial metagenomic signature, we also assessed the association between viral composition and disease severity, and found a different trend presented in the viral component. While bacterial abundances are reduced in critical patients as compared to non-critical patients, viral abundances in critical patients are higher than in non-critical patients (**Fig. 2E and S7**). Also different from the bacterial signature, the viral species richness does not significantly change, but the Shannon diversity of viral species in critical patients is significantly lower (**Fig. 2E**). This effect is partly contributed by a greater abundance of SARS-Cov-2 sequences, which could then lead to a lower Shannon Index as signals from low abundance viruses could be drowned out. Intriguingly, several other viral families display increased abundance in critical patients as compared to non-critical (**Fig. 2F**), including many dsDNA viruses that are known to establish latency such as herpesviruses and papillomaviruses. One speculation is that the correlation of abundance of these viral families with disease severity could be due to reactivation of latent viruses under immunosuppressive medication, changes in host immune activity, or direct SARS-CoV-2 activity^15–17^. The effect of age that we observed for bacteria is less pronounced in viruses (**Fig. S8**), and is only observed for some viral families. Additional sampling and deeper viral sequencing, as well as systematic experimental designs are needed to further investigate these phenomena.

For severe viral pneumonia, co-infections can greatly affect patient outcomes^18–20^. One recent study has showed that 50% of patients with COVID-19 who have died in this pandemic had secondary bacterial infections^21^. By surveying the metagenomic landscape of these samples, we observed several patient samples with exceptionally high abundance of known pathogens, which could indicate a co-infection with SARS-Cov-2 in those patients. We found ten cases of *S. aureus* and nine cases of *C. albicans* co-infections, and the rate of co-infection for both pathogens is generally correlated with disease severity (**Fig. 2G**).

### MINERVA achieves better SARS-CoV-2 genome coverage compared to conventional dsDL strategies

In both SHERRY and dsDL data, we detected low yet significant levels of SARS-CoV-2 sequences. The viral ratio is between 10^-7^ and 10^-1^. It is worth noting that the SARS-CoV-2 sequence ratio is higher in SHERRY data than in dsDL data (**Fig. 3A and 3B**), suggesting that SHERRY libraries capture more SARS-CoV-2 sequences. Though SARS-CoV-2 genome coverage and depth was not high in SHERRY results due to low viral ratio and low sequencing depth, performing MINERVA subsequently can enrich the SARS-CoV-2 sequence ratio up to 10,000-fold (**Fig. 3C and S9**). As a result, MINERVA gives more complete and deeper coverage of SARS-CoV-2 genomes (**Fig. 3D and 3E**), despite sequencing dsDL libraries to two orders of magnitude more depth (**Fig. S3**).

**Figure 3.**
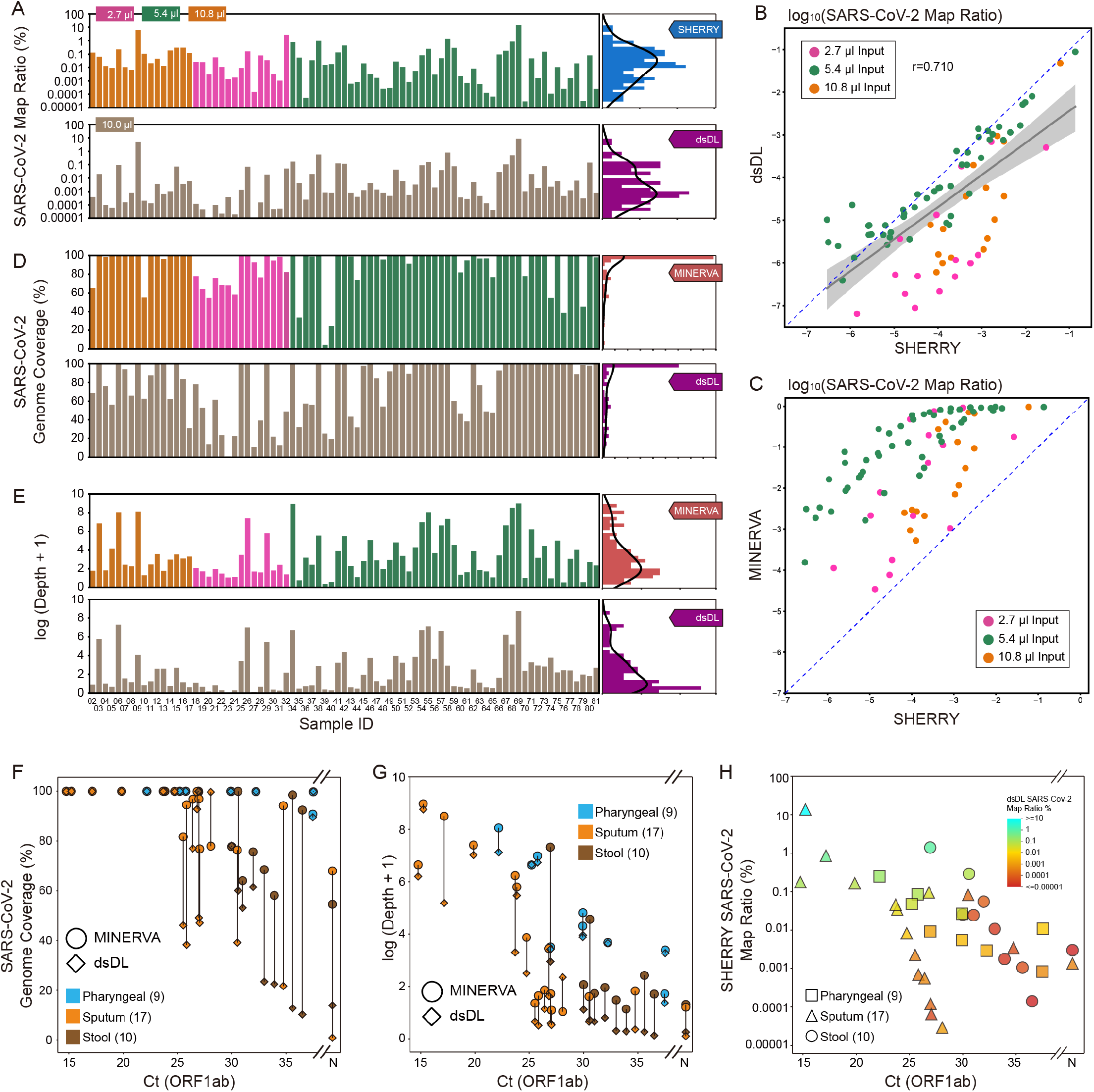
Direct comparison between sequencing libraries constructed from MINERVA and conventional dsDL strategies. (A) SARS-CoV-2 mapping ratio statistics of SHERRY and dsDL libraries. (B) Comparison of SARS-CoV-2 mapping ratios between SHERRY and dsDL libraries. (C) Comparison of SARS-CoV-2 mapping ratios between SHERRY and MINERVA libraries. (D and E) SARS-CoV-2 genome coverage and depth statistics of MINERVA and dsDL libraries. (F and G) Comparison of SARS-CoV-2 sequencing results between MINERVA and dsDL libraries. (H) Metagenomic sequencing and qPCR result features of samples with poor SARS-CoV-2 genome coverage.

The superior quality of MINERVA data became clearer when we included clinical RT-qPCR results. Both dsDL and MINERVA libraries detect SARS-CoV-2 sequences for samples with various Ct values, but MINERVA produced more complete and deeper genome coverage than dsDL methods (**Fig. 3F and 3G**), and this advantage is more pronounced for low viral load samples, including two samples with negative qPCR results, and stool samples. By studying the relationship between SARS-CoV-2 qPCR results and read ratio, we identified two groups of samples that resulted in low SARS-CoV-2 genome coverage when processed using dsDL (**Fig. 3H**). The first group had low SARS-CoV-2 read ratio, which prohibited the acquisition of enough SARS-CoV-2 sequencing reads.

The second group, which included most stool samples, had relatively high SARS-CoV-2 Ct values and read ratio, suggesting these samples had low total nucleic acid amount. Since dsDL approaches are less sensitive and require more input, this may explain why MINERVA outperforms dsDL most evidently in stool samples.

### MINERVA can facilitate multiple facets of COVID-19 research

As a novel virus, little is known about the evolutionary features of SARS-CoV-2. Using 143 samples, we constructed a SARS-CoV-2 mutational profile (**Fig. 4A**), which was distinct from the Guangdong profile^22^. A few mutation sites, including the two linked to S and L strains^23^, were found in multiple samples. Aided by the deep genome coverage in MIVERVA data, we not only detected strong linkage between position 8,782 and 28,144, but also observed high concordance of allele frequencies between these two positions. Furthermore, we detected strong linkage and high allele frequency concordance among four other positions, 241, 3,037, 14,408 and 23,403. Such allele frequency information offers additional layers of evidence supporting co-evolution of positions within the SARS-CoV-2 genome, in two distinct groups of samples. It is worth noting that in some samples, not all linked alleles are simultaneously detected, due to low coverage at some positions in those samples; these alleles can indeed be observed at low coverage in the raw data for these samples, but is missing from the post-processing data as they do not pass the stringent quality filtering steps. Nonetheless, the linkage was established by observing such linkage over many samples.

**Figure 4.**
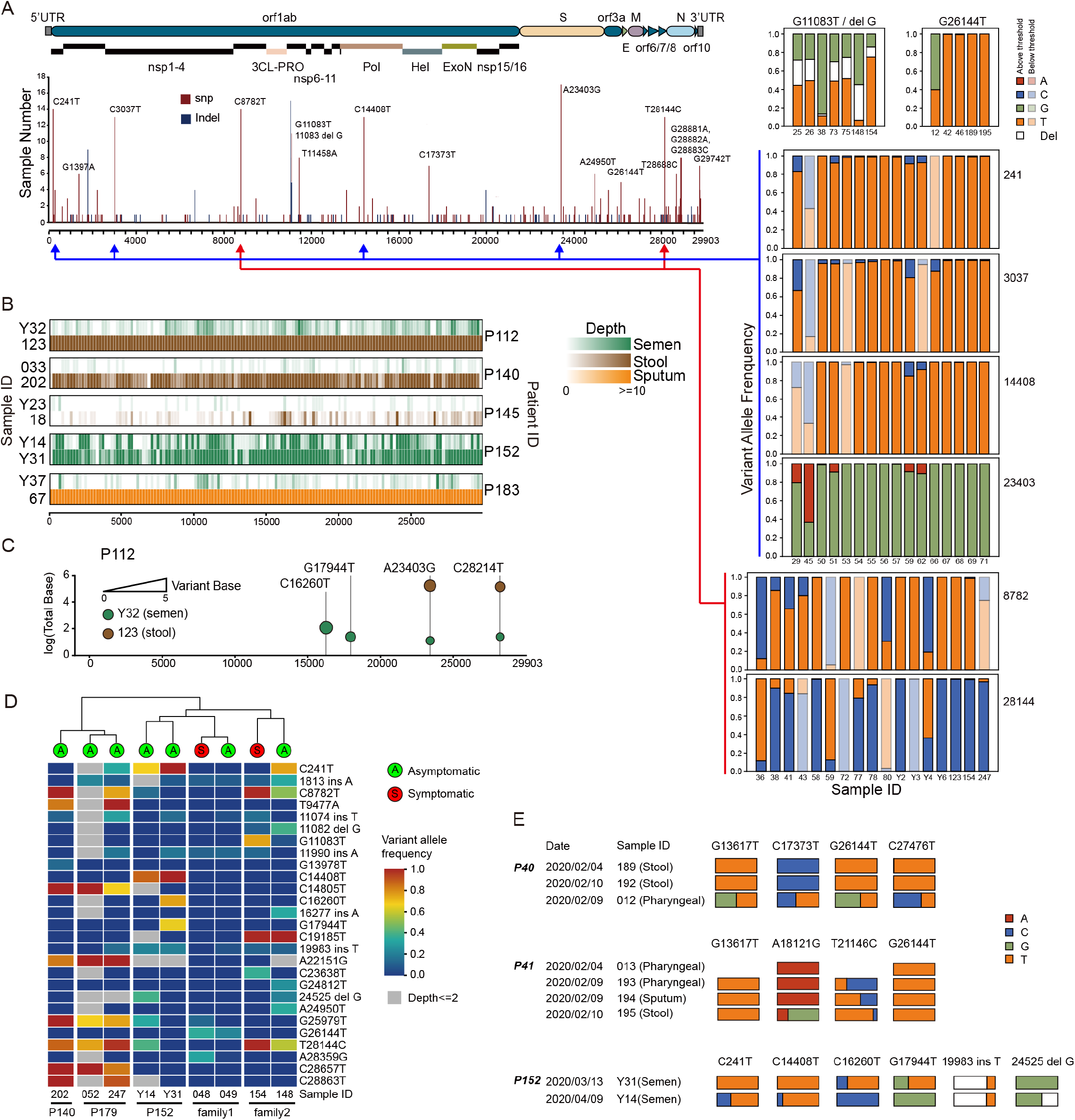
MINERVA could facilitate COVID-19 and SARS-CoV-2 research through accurate and sensitive identification of viral mutations. (A) SARS-CoV-2 mutation profile obtained from 101 samples. (B) SARS-CoV-2 genome coverage of semen samples and other types of samples from the same patients. (C) SARS-CoV-2 mutations in semen and stool samples from Patient 112. (D) SARS-CoV-2 mutation profiles of asymptomatic patients and their infected family members. (E) Longitudinal SARS-CoV-2 mutation analysis of individual patients.

Several studies have examined the distribution of SARS-CoV-2 across different organs and tissues^7^. However, the presence of SARS-CoV-2 in the reproductive system is still under debate^24^,^25^. Aided by the high sensitivity of MINERVA, we detected a high abundance of SARS-CoV-2 sequences in semen samples from COVID-19 patients (**Fig. 4B**); several semen samples also had high SARS-CoV-2 genome coverage. SARS-CoV-2 SNV analysis demonstrated high similarity between semen and non-semen results (**Fig. 4C**).

Apart from its high infectiousness, the containment of SARS-CoV-2 transmission is challenging due to the existence of asymptomatic infected individuals^26^. Though RT-qPCR can be used to identify these individuals, elucidation of the chain of transmission requires the complete SARS-CoV-2 sequences. To evaluate the performance of MINERVA for tracking SARS-CoV-2 transmission, we sequenced the samples of several asymptomatic individuals and their infected family members. SARS-CoV-2 SNV analysis revealed that asymptomatic individuals each harbor viral sequences with unique signatures, however, these individuals are clustered by the viral SNV signature with their respective family members rather than other asymptomatic individuals, which indicates that viral SNVs within infected families are similar to each other and unique from other families (**Fig. 4D**). Summarily, despite the asymptomatic phenotype of some infected individuals, the viral SNV signature generated by MINERVA can be used to accurately place these individuals in the chain of transmission, enabling better epidemiological tracking.

Recent studies have identified genetic variations of SARS-CoV-2 and raised the possibility that multiple variants could co-exist in the same host individual. The intra-host SNVs (iSNVs) detected in many samples (**Fig. 4A**) suggest that SARS-CoV-2 is rapidly evolving within the hosts^2^. Through longitudinal sampling, we confirmed that iSNVs were generally relatively stable across time and body sites (**Fig. S10**), but found that some patients harbored greater variations in iSNVs (**Fig. 4E**). For P40 and P41, iSNVs were stable within the same sample type across time, but varied across different sample types. Comparing the two semen samples from Patient 152, changes in iSNV were clearly observed. These results support the co-existence of multiple SARS-CoV-2 variants in the same individual, and further investigation is warranted to understand this phenomenon.

In summary, MINERVA effectively converts metagenomes and SARS-CoV-2 sequences into sequencing libraries with a simple and quick experimental pipeline, and subsequent target enrichment can further improve SARS-CoV-2 genome coverage and genetic variation detection. MINERVA can facilitate the study of SARS-CoV-2 genetics, and be easily implemented to fight future RNA pathogen outbreaks.

## Discussion

As of today, our knowledge of SARS-CoV-2 is still preliminary and much of it extrapolated from past studies of other beta coronaviruses such as SARS and MERS. However, the epidemiology, physiology, and biology of COVID-19 are evidently unique^27^. To speed up our investigation of this virus and the disease it causes, a practical protocol for viral genome research of clinical samples is urgently needed. Currently, methods for transforming clinical samples into sequencing libraries are laborious and painstaking, while clinical personnel at the frontlines are already strained for time and energy. MINERVA minimizes the need for expert technique and hands-on operation; we believe it will be pivotal in accelerating clinical research of SARS-CoV-2.

Recent evolutionary tracing studies suggest the emergence of multiple novel, evolved subtypes of SARS-CoV-2^28^, including the S/L-subtypes^23^ and the A/B/C-variants^29^. New variants will likely continue to emerge as the virus mutates, and to uncover them requires deep, complete coverage of viral genomes from a large number of patients. With the existence of asymptomatic carriers^26^ and possible recurrent infections in the same individual^30^, longitudinal re-sampling of patients is also important to uncover intra-host viral heterogeneity, but as viral load decreases with time^31^, the sensitivity of the sample processing method becomes critical. These studies all require processing large volumes of clinical samples with a highly robust and scalable method that does not compromise on sensitivity. We have demonstrated that MINERVA libraries from clinical samples can generate deep and complete coverage of SARS-CoV-2 genomes that can be used for evolutionary tracing and variant characterization research. Furthermore, the high sensitivity, high coverage, and high depth of the SARS-CoV-2 viral genomes obtained by MINERVA can reveal unique viral SNV signatures in each patient, even if they are asymptomatic. We showed that these viral SNVs allows families of infected individuals to be co-clustered, but are unique between families, which enables each individual to be accurately placed in the chain of transmission. As MINERVA is easily scalable and implementable in a clinical lab setting, it can serve as a robust strategy for timely and critical epidemiological tracking and monitoring during a pandemic.

It is well-established now that SARS-CoV-2 can infect multiple organ systems, tissue compartments, and cell types^2^,^7^,^32^,^33^. In our profiling of COVID-19 clinical samples from multiple body sites of the same patient, we found that the viral load and viral subtypes vary across different body sites, possibly affected by interactions between microbial and other viral species as well as overall metagenomic diversity present in different microenvironments of each body site. The effects of metagenomic diversity and inter-compartment heterogeneity on SARS-CoV-2 biology and COVID-19 symptom severity are also not understood. In particular, it is difficult to obtain high-quality unbiased metagenomic using conventional library construction methods from low-quantity samples, as well as samples such as stool in which bacteria dominate the metagenomes, as conventional methods are not sufficiently sensitive. The versatility of MINERVA as a two-part protocol integrating a tailored SHERRY and post-library virus enrichment provides flexibility for sample processing that uses one standard sample pipeline for both highly sensitive metagenomic analysis and targeted deep sequencing of specific transcripts. Using MINERVA, we have demonstrated the first large scale profiling of metagenomic composition of different body sites in the context of COVID-19. One pre-print study investigated the relationship between gut microbes and COVID-19 severity, and purportedly found links between gut microbe composition with blood proteomic biomarkers that predict symptom severity^34^, however, there is no discussion of metagenomic composition of other body sites. As we show here with MINERVA data from a wide range of sample types, there are large body site-specific differences, and our data suggests that microbial and viral metagenomic composition in pharyngeal swab samples also significantly correlates with disease severity. The metagenomic profile of these other body sites that are arguably more directly involved in the viral infection, have not been reported or investigated elsewhere. Using MINERVA we highlight several new directions of clinical and basic research, and with further investigation, these could shed light on the complex interactions between SARS-CoV-2 pathology, host microbial communities, host immunity, and disease progression. We also showed that MINERVA metagenomic profiles can identify potential co-infections of bacteria, fungi, and other viruses, which is challenging to do with conventional approaches. In our samples, we found a co-infection rate of ~ 20% (16/79 patients), which is higher than the rate reported by one secondary study of COVID-19 co-infections^35^. In this secondary study, although they found 8% of patients experiencing bacterial/fungal co-infection, the rate of broad spectrum antibiotic use for COVID-19 patients is much higher (72%). It is well-known that co-infections in severe pneumonia can greatly affect patient outcomes^19^,^20^, and it is estimated that 50% of patients with COVID-19 who have died in this pandemic had secondary bacterial infections^21^. Our result shows the utility of MINERVA in identifying non-viral co-infections, and further primary studies using MINERVA could help to elucidate true co-infection rates to guide better strategies for antibiotic use.

MINERVA was not created to be a rapid diagnostic assay; rather, we hope its ease-of-use, versatility, scalability, sensitivity, and cost-effectiveness will drive adoption of routine sequencing of COVID-19 clinical samples, and thereby facilitate multiple areas of much-needed SARS-CoV-2 and COVID-19 research for clinicians and researchers.

## Author contributions

C.C., Y.C., X.S.X., H.Z., Y.H. and J.W. conceived the project; J.L., P.D., Q.L. and C.S. conducted experiments; C.C., L.D., Q.J., J.L., Y.H. and J.W. analyzed the data; C.C., J.L., L.D., Q.J., A.R.W., Y.H. and J.W. wrote the manuscript with the help from all other authors.

## Acknowledgement

We thank Ms. Chenyang Geng and BIOPIC sequencing platform at Peking University for the assistance of high-throughput sequencing experiments, and Ms. Amelia Huang for the assistance of figure preparation. This work was supported by National Natural Science Foundation of China (21675098, 21927802, 21525521), Ministry of Science and Technology of China (2018YFA0800200, 2018YFA0108100, 2018YFC1002300), 2018 Beijing Brain Initiation (Z181100001518004), Beijing Advanced Innovation Center for Structural Biology, Beijing Advanced Innovation Center for Genomics, HKUST’s start-up and initiation grants (Hong Kong University Grants Committee), Hong Kong Research Grants Council Theme-based Research Scheme (RGC TBRS T12-704/16R-2) and Collaborative Research Fund (RGC CRF C6002-17G), Hong Kong RGC Early Career Support Scheme (RGC ECS 26101016), Hong Kong Epigenomics Project (LKCCFL18SC01-E), and HKUST BDBI Labs.

---

**Figure S1**. Comparison of workflow between MINERVA and the conventional dsDL strategy.

**Figure S2**. Optimization of SHERRY protocol. (A-C) Effect of N10 primer during reserve transcription and Tn5 amount on detected gene number, ribosomal rate and insert size. (D-F) Effect of N10 primer during reserve transcription and Tn5 amount on gene body coverage evenness.

**Figure S3**. Amount of sequencing data for different libraries.

**Figure S4**. Comparison between SHERRY and dsDL libraries on total viral ratio (A), total fungal ratio (B), total bacterial ratio (C), and bacterial entropy (D).

**Figure S5**. PCA analysis of viral and fungal compositions in different sample types. (A) Viral family composition in all samples. The viral composition of faeces samples was distinct from oral samples. (B) Fungal family composition in all samples. There is no major difference among different sample types and stages. While *Candida* can be detected with high level in certain patients.

**Figure S6**. Bacterial composition by severity in different sample types and age groups. (A) Bacterial composition by severity in different sample types. The severity stage is highly related to age. The bacterial ratio and species richness were significantly lower in critical pharyngeal samples (Kruskal-Wallis test and Dunn’s post-hoc test). This was not observed in sputum samples may be because of the small sample size. (In pharyngeal samples, mild=15, moderate=19, severe=8, critical=18; while in sputum samples, mild=8, moderate=34, severe=8, critical=1). (B) Comparison of bacterial ratio and within-subject diversity between non-critical (n=8) and critical patients (n=16) in old group (>=60 years old) to avoid the bias from age. The bacterial ratio and species richness were lower in critical patients while the Shannon index is higher (Mann-Whitney U test). (C) Comparison of bacterial ratio and within-subject diversity between critical (n=2) and non-critical patients (n=34) in young group (<60 years old). The bacterial ratio and species richness were lower in critical patients while the Shannon index is higher.

**Figure S7**. Viral composition by severity in different sample types. The viral ratio was significantly lower in critical patients compared to other patients in pharyngeal samples (Kruskal-Wallis test and Dunn’s post-hoc test).

**Figure S8**. Bacterial composition by severity in different age groups. (A) Comparison of relative abundance of SARS-Cov-2 and Coronovaridae between non-critical (n=8) and critical (n=16) old (>=60) patients in pharyngeal samples. Critical patients have higher level of SARS-Cov-2 and Coronaviridae (Mann-Whitney U test). (B) Comparison of relative abundance of SARS-Cov-2 and Cronovaridae between non-critical (n=34) and critical (n=2) young (<60) patients in pharyngeal samples. Critical patients have higher level of SARS-Cov-2 and Coronaviridae (Mann-Whitney U test). (C) Comparison of relative abundance of Herpesviridae, Papillomaviridae and Poxviridae between non-critical (n=8) and critical (n=16) old (>=60) patients in pharyngeal samples. The abundance of these three viral families was higher in Critical patients (Mann-Whitney U test). (D) Comparison of relative abundance of Herpesviridae, Papillomaviridae and Poxviridae between non-critical (n=8) and critical (n=16) young (<60) patients in pharyngeal samples. The abundance of these three viral families was higher in Critical patients (Mann-Whitney U test).

**Figure S9**. SARS-CoV-2 genome sequencing results of SHERRY and MINERVA libraries. (A) SARS-CoV-2 mapping ratio statistics of MINERVA libraries. (B and C) SARS-CoV-2 genome coverage and depth statistics of SHERRY libraries.

**Figure S10**. Longitudinal SARS-CoV-2 mutation analysis of individual patients.

## Material and Methods

### Ethics approval

This study was approved by the Ethics Committee of Beijing Ditan Hospital, Capital Medical University (No. KT2020-006-01).

### Optimization of SHERRY protocol

We used the total RNA extracted from 3T3 cells to optimize experimental protocols. RNA extraction was performed using RNeasy Mini Kit (Qiagen, Cat.No.74104). DNA was then removed through DNase I (NEB, Cat.No.M0303) digestion. The resulting total RNA was concentrated by RNA Clean & Concentrator-5 kit (Zymo Research, Cat R1015), and its quality was assessed by the Fragment Analyzer Automated CE System (AATI). Its quantification was done by Qubit 2.0 (Invitrogen). To optimize the SHERRY protocol, different amount of random decamer (N10) (0, 10, or 100 pmol) was used to set up reverse transcription reactions. Titration of Tn5 transposome (0.2, 0.5, or 1.0 μl Vazyme V50; 0.05 or 0.25 μl home-made pTXB1) was performed in tagmentation procedure. In all tests, 10 ng 3T3 total RNA was used, and all reagents except for N10 or Tn5 transposome remain unchanged. All libraries were sequenced on Illumina NextSeq 500 with 2×75 paired-end mode. Clean data was aligned to GRCm38 genome and known transcript annotation using Tophat2 v2.1.1. Ribosome-removed aligned reads were proceeded to calculate FPKM by Cufflinks v2.2.1 and gene body coverage by RSeQC v.2.6.4.

### Patients and clinical samples

From January 23, 2020 to April 20, 2020, 91 patients were enrolled in this study according to the 7th guideline for the diagnosis and treatment of COVID-19 from the National Health Commission of the People’s Republic of China. All patients, diagnosed with COVID-19, were hospitalized in Beijing Ditan Hospital and classified into four severity degrees, mild, moderate, severe, and critical illness, according to the guideline. We collected 143 samples (60 pharyngeal swabs, 52 sputum samples, 25 stool samples, and 6 semen samples) from these patients.

### RNA extraction and rRNA removal

For all the clinical samples, nucleic acids extraction was performed in a BSL-3 laboratory. Samples were deactivated by heating at 56°C for 30 min before extraction. Total RNA was extracted using QIAamp Viral RNA Mini Kit (Qiagen) following the manufacturer’s instructions. In most samples (79 out of 85) we specifically omitted the use of carrier RNA due to its interference on the most prevalent sample preparation protocols for high-throughput sequencing. After nucleic acids extraction, rRNA was removed by rDNA probe hybridization and RNase H digestion, followed by DNA removal through DNase I digestion, using MGIEasy rRNA removal kit (BGI, Shenzhen, China). The final elution volume was 12-20 μl for each sample. For carrier RNA removal tests, 1.7 μg polyA carrier RNA was spiked into 18 μl of elute from QIAamp Viral RNA Mini Kit. To remove the carrier RNA from these spike-in samples and other samples extracted with carrier RNA, 2 μg poly(T) 59-mer (T59) oligo was added during the rDNA hybridization step.

### dsDL Metagenomic RNA library construction and sequencing

The libraries were constructed using MGIEasy reagents (BGI, China) following manufacture’s instruction. The purified RNA, after rRNA depletion and DNA digestion, underwent reverse transcription, second strand synthesis, and sequencing adaptor ligation. After PCR amplification, DNA was denatured and circularized before being sequenced on DNBSEQ-T7 sequencers (BGI, China).

### MINERVA library preparation

Totally, 2.7 μl RNA from rRNA and DNA removal reaction was used for standard SHERRY reverse transcription, with the following modifications: 1) 10 pmol random decamer (N10) was added to improve coverage; 2) initial concentrations of dNTPs and oligo-dT (T30VN) were increased to 25 mM and 100 μM, respectively. For 5.4 μl and 10.8 μl input, the entire reaction was simply scaled up 2 and 4 folds, respectively. The RNA/DNA hybrid was tagmented in TD reaction buffer (10 mM Tris-Cl pH 7.6, 5 mM MgCl_2_, 10% DMF) supplemented with 3.4% PEG8000 (VWR Life Science, Cat.No.97061), 1 mM ATP (NEB, Cat.No. P0756), and 1U/μl RNase inhibitor (TaKaRa, Cat.No. 2313B). The reaction was incubated at 55°C for 30 min. 20 μl tagmentation product was mixed with 20.4 μl Q5 High-Fidelity 2X Master Mix (NEB, Cat.No. M0492L), 0.4 μl SuperScript II reverse transcriptase, and incubated at 42°C for 15 min to fill the gaps, followed by 70°C for 15 min to inactivate SuperScript II reverse transcriptase. Then index PCR was performed by adding 4 μl 10 μM unique dual index primers and 4 μl Q5 High-Fidelity 2X Master Mix, with the following thermo profile: 98°C 30 s, 18 cycles of [98°C 20 s, 60°C 20 s, 72°C 2 min], 72°C 5 min. The PCR product was then purified with 0.8x VAHTS DNA Clean Beads (Vazyme, Cat. No. N411). These SHERRY libraries were sequenced on Illumina NextSeq 500 with 2×75 paired-end mode for metagenomic analysis.

For preparing MINERVA libraries through SARS-CoV-2 enrichment, 1μl SHERRY metagenomic library was first quantified with N gene using quantitative PCR (F: GGGGAACTTCTCCTGCTAGAAT, R: CAGACATTTTGCTCTCAAGCTG) after 1:200 dilution, then multiple libraries were pooled together based on qPCR results and processed with TargetSeq One Cov Kit (iGeneTech, Cat.No.502002-V1) following manufacturer’s instruction. The iGeneTech Blocker was replaced by the IDT xGen Universal Blockers (NXT). These MINERVA libraries were sequenced on Illumina NextSeq 500 with 2×75 paired-end mode for deep SARS-CoV-2 analysis.

### Data processing

For metagenomic RNA-seq data, raw reads were quality controlled using BBmap (version 38.68) and mapped to the human genome reference (GRCh38) using STAR (version 2.6.1d) with default parameters. All unmapped reads were collected using samtools (version 1.3) for microbial taxonomy assignment by Centrifuge (version 1.0.4). Custom reference was built from all complete bacterial, viral and any assembled fungal genomes downloaded from NCBI RefSeq database (viral and fungal genomes were downloaded on February 4th, 2020, and bacterial genomes were downloaded on November 14th, 2018). There were 11,174 bacterial, 8,997 viral, and 308 fungal genomes respectively. Bacterial Shannon diversity (entropy) was calculated at species level, and the species abundance was measured based on total reads assigned at the specific clade normalized by genome size and sequencing depth. Bacterial genus composition was analyzed based on reads proportion directly assigned by Centrifuge. For dsDL sequencing data, sub-sampling was performed for each sample to obtain ~12M paired-end nonhuman reads, which is the median of SHERRY datasets. Same workflow was performed as described above for the removal of human reads and microbial taxonomy assignment.

For SARS-CoV-2 genome analysis, raw reads were trimmed to remove sequencing adaptors and low-quality bases with Cutadapt v1.15. BWA 0.7.15-r1140 was used to align reads to the SARS-CoV-2 reference genome (NC_045512.2). Then we removed duplicates from the primary alignment with Picard Tools v2.17.6. We used mpileup function in samtools v1.10 to call SNP and InDel with parameter -C 50 -Q 30 -q 15 -E -d 0. We called mutation if the depth >=10 and strand bias >0.25. The strand bias is defined as the value that minimum of positive strand depth and negative strand depth divided by the maximum.

### Data deposition

The sequencing data generated during this study have been uploaded to Genome Sequencing Archive (PRJCA002533). However, due to ethical concerns, access to the datasets is only available from the corresponding author on reasonable request.

