## Supplementary material for "MINERVA: A facile strategy for SARS-CoV-2 whole genome deep sequencing of clinical samples": SI

Supplementary Figure S1

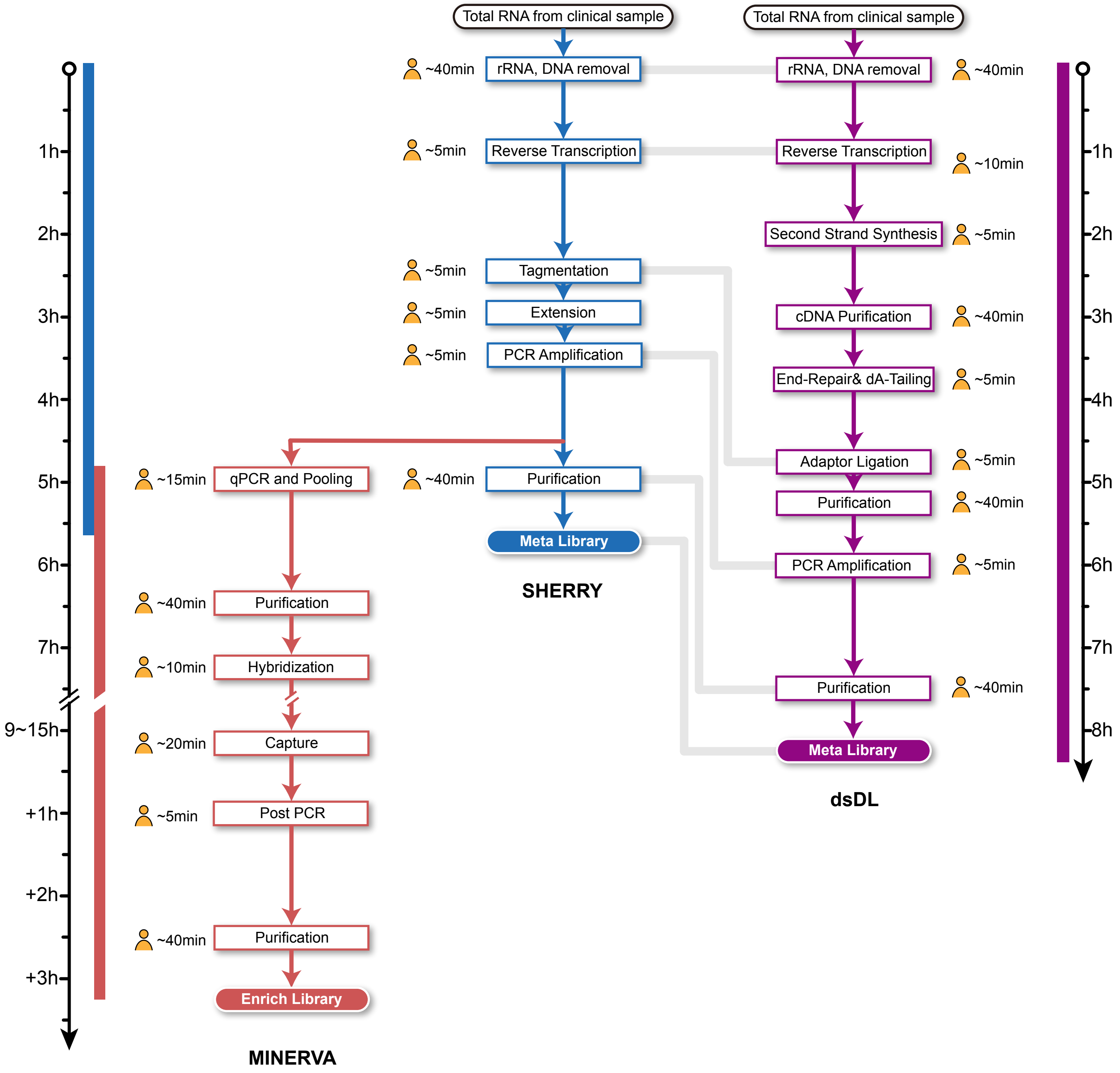

Supplementary Figure S2

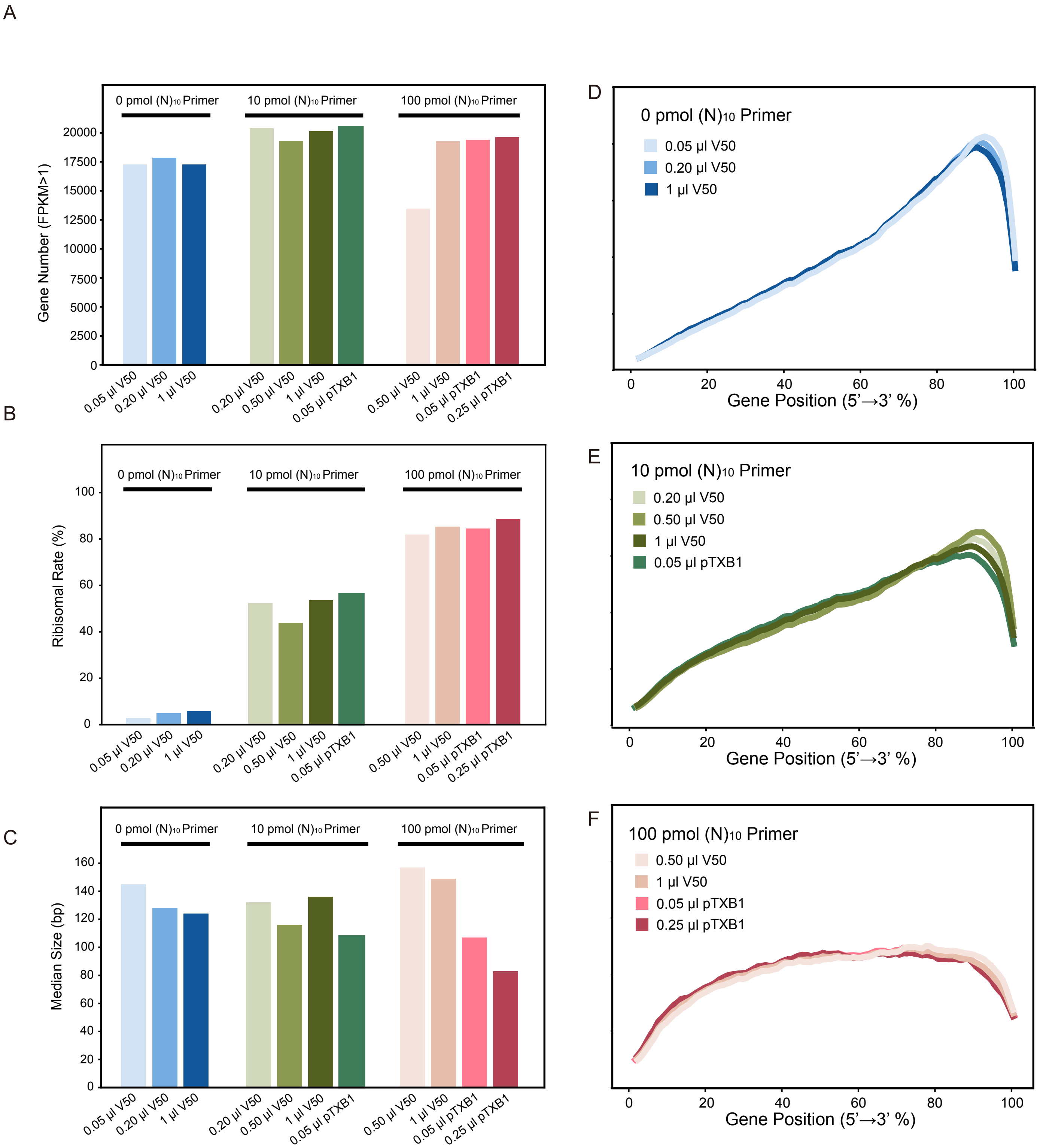

Supplementary Figure S3

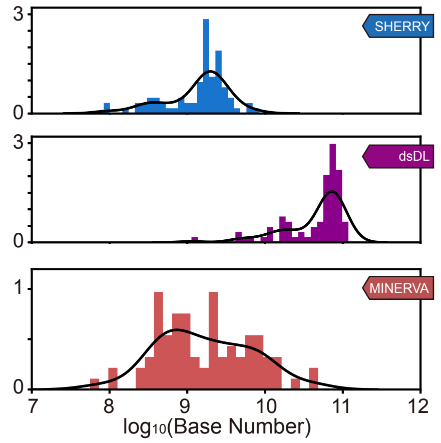

Supplementary Figure S4

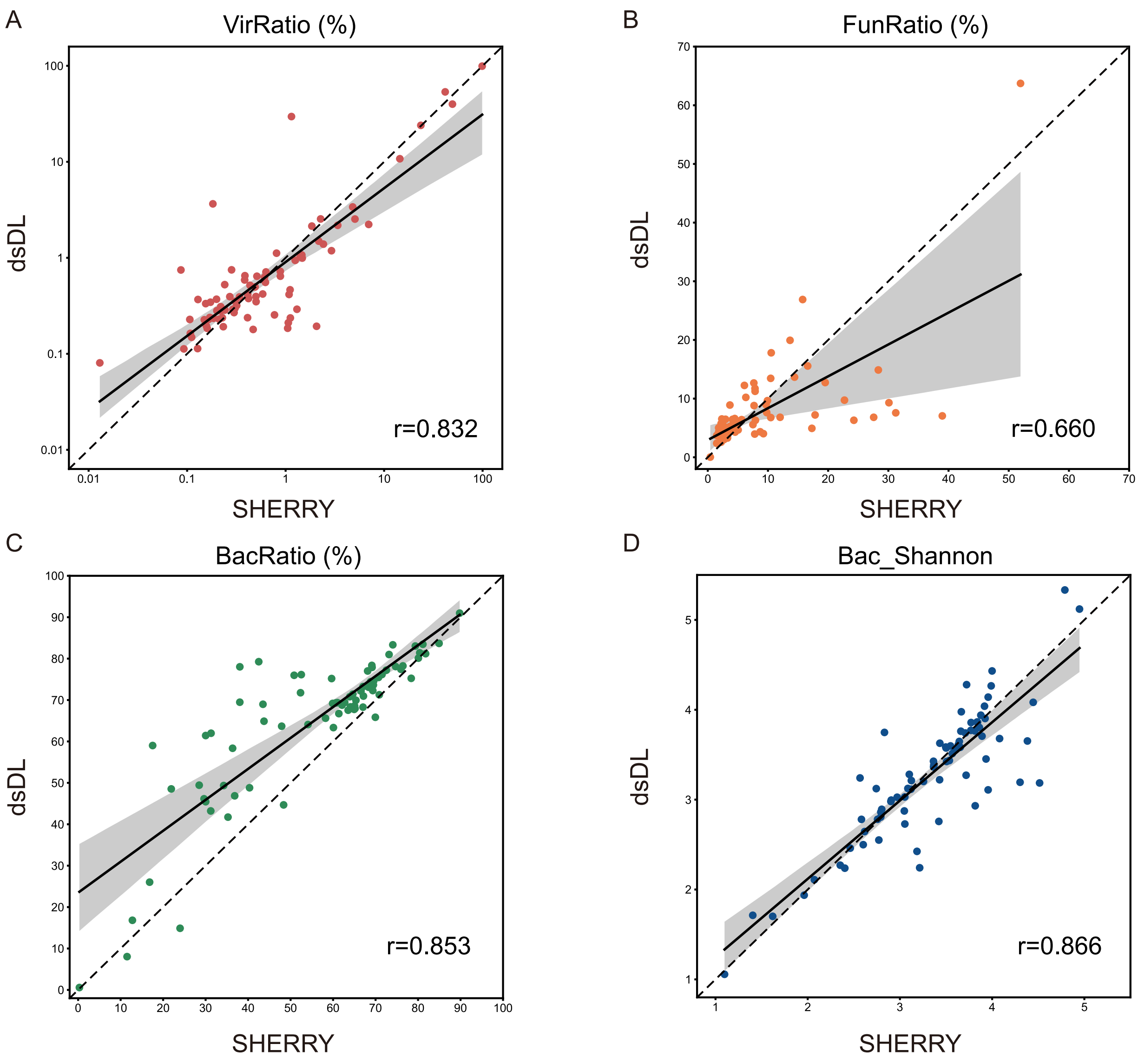

Supplementary Figure S5

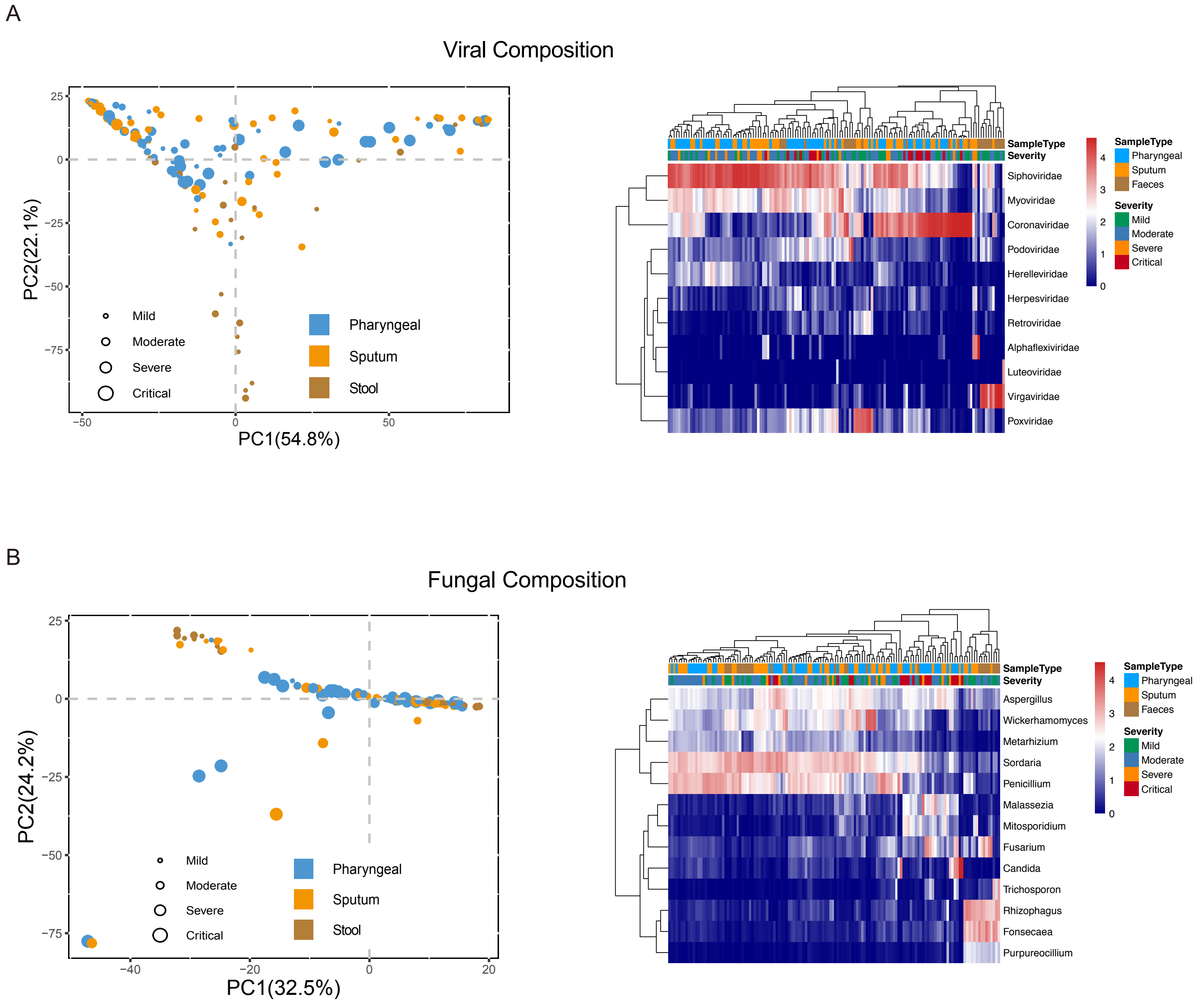

Supplementary Figure S6

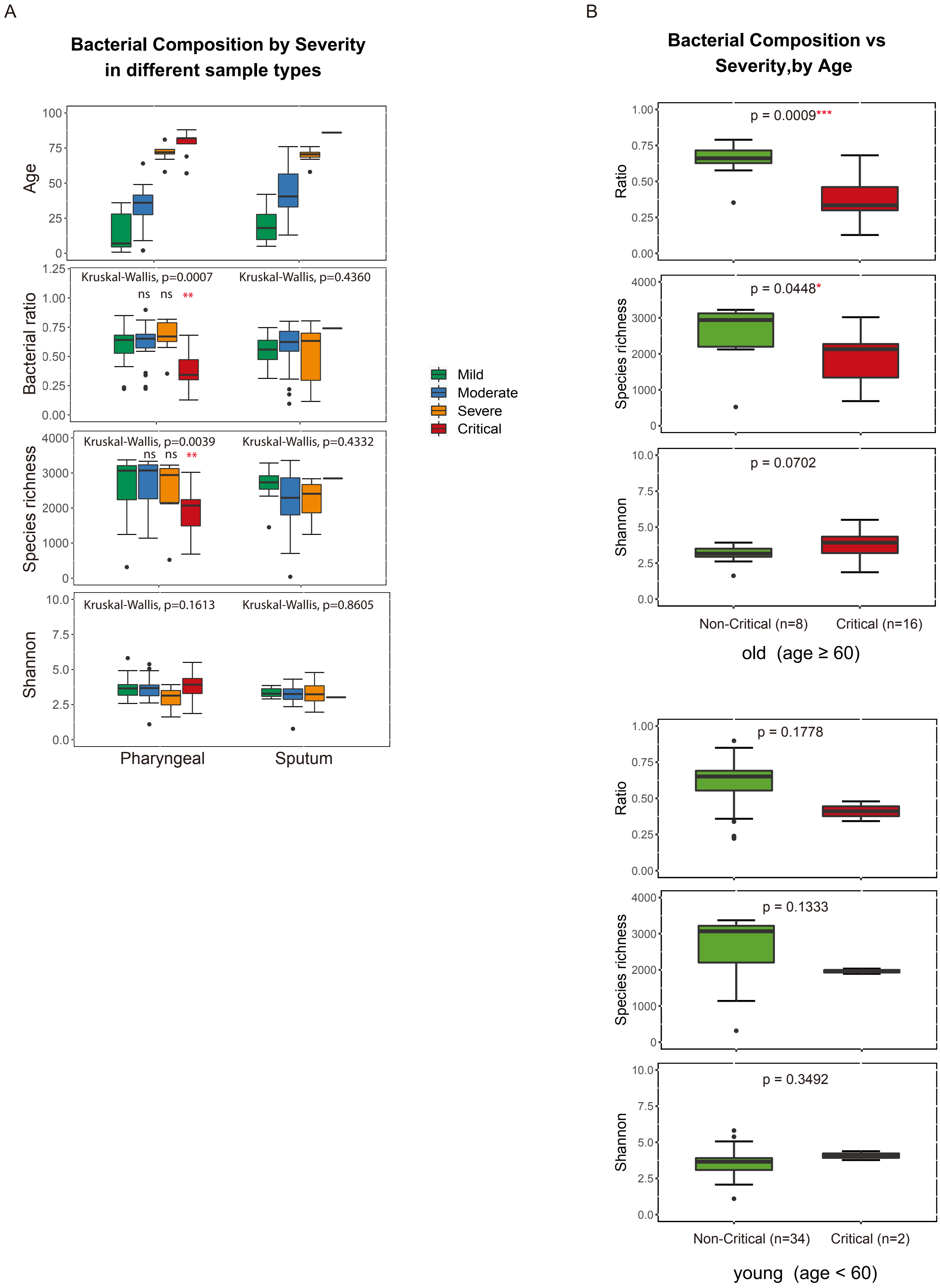

Supplementary Figure S7

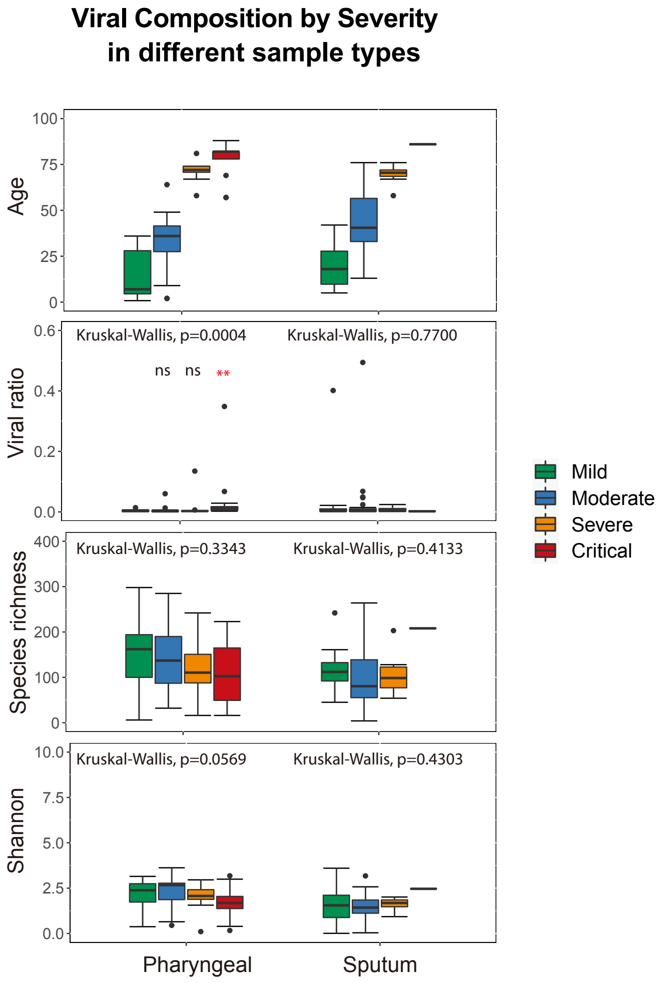

Supplementary Figure S8

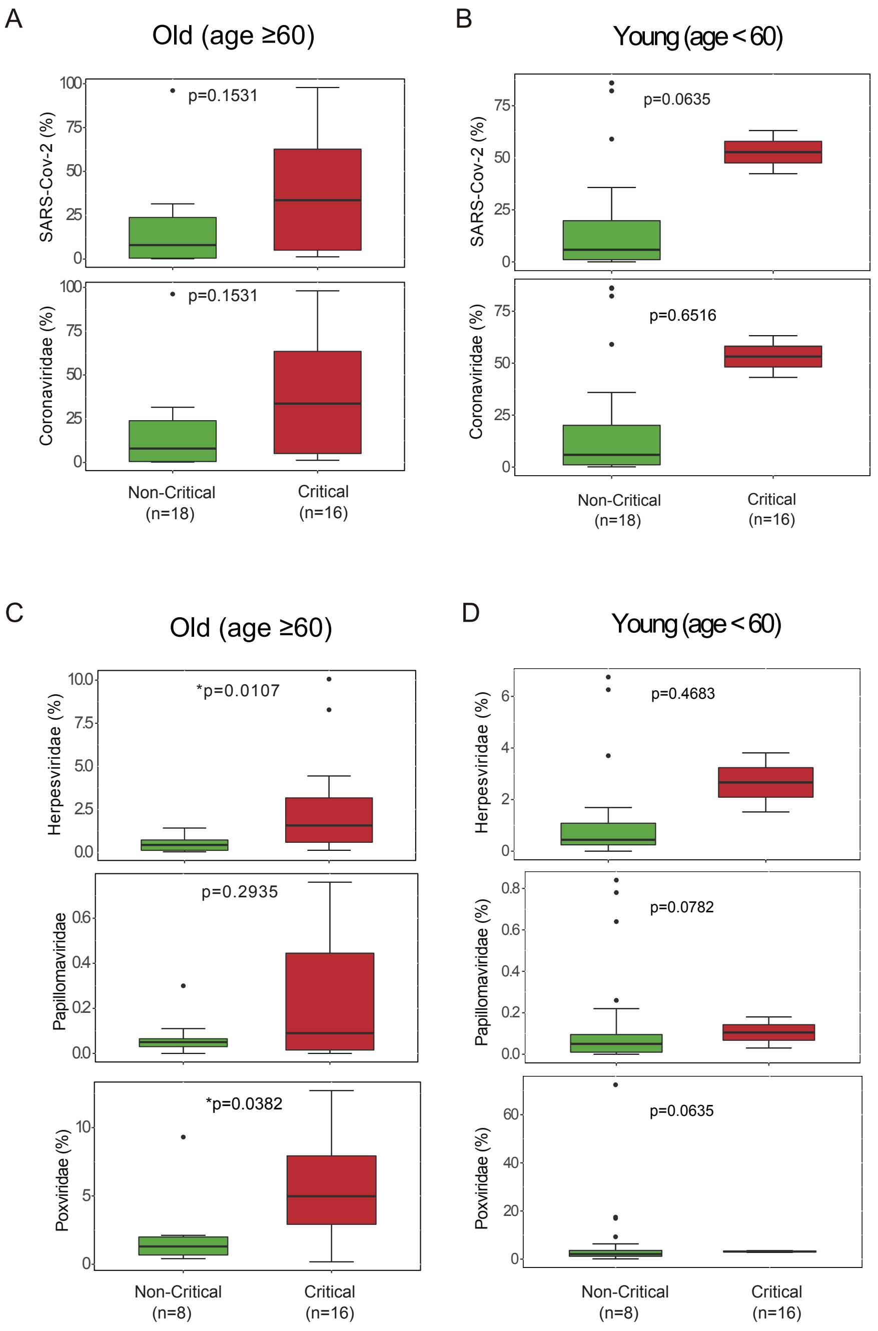

Supplementary Figure S9

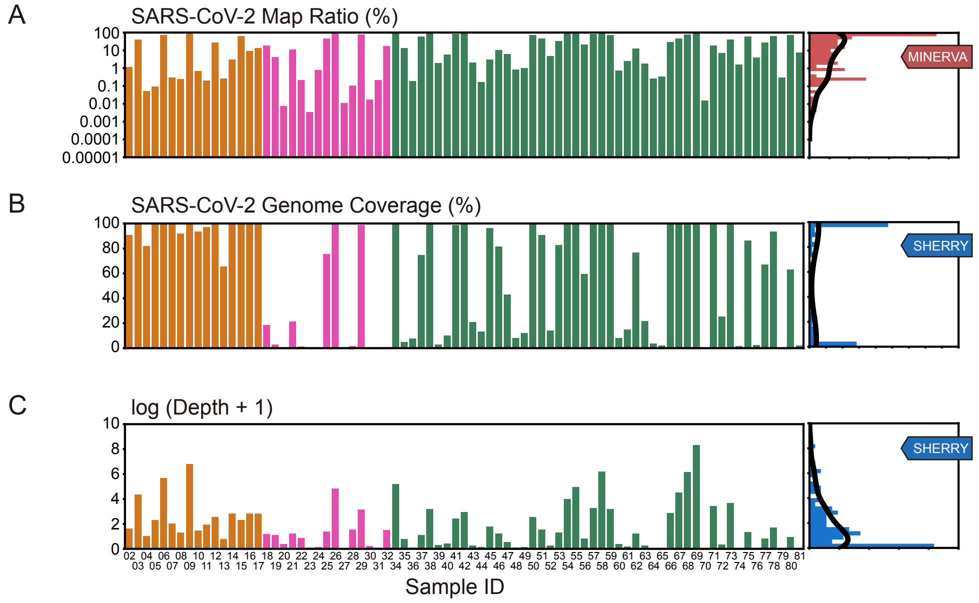

Supplementary Figure S10

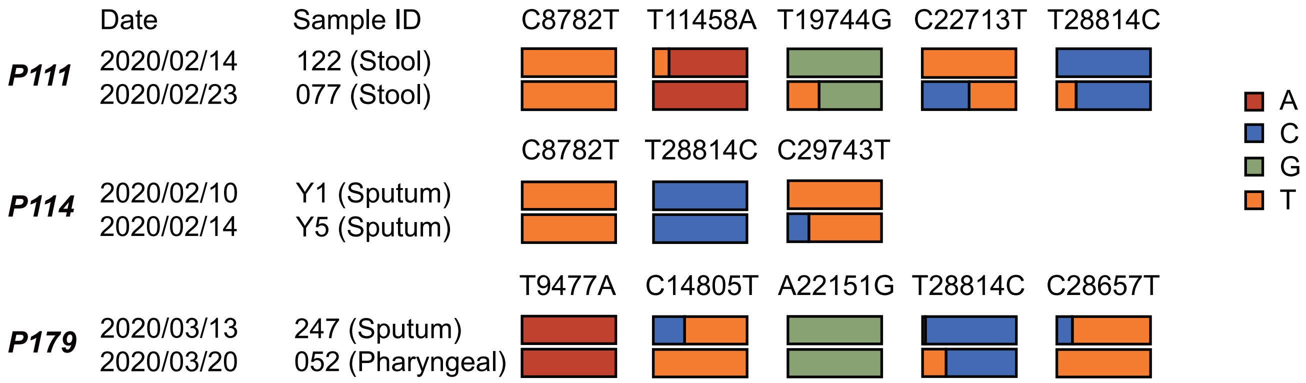

| | MINERVA | | | | | SHERRY | | | | | dsDL | | | |
|---|---|---|---|---|---|---|---|---|---|---|---|---|---|---|
| Sample ID | Input (μl) | Total Reads | Mapping Ratio | Dedup Coverage | Dedup Depth | Input (μl) | Total Reads | Mapping Ratio | Dedup Coverage | Dedup Depth | Input (μl) | Mapping Ratio | Dedup Coverage | Dedup Depth |
| 02 | 10.8 | 8,195,484 | 1.1638% | 65.03% | 4.9783 | 10.8 | 5,094,918 | 0.1307% | 90.87% | 4.0136 | 10.0 | 0.0004% | 68.96% | 1.4060 |
| 03 | 10.8 | 207,709,272 | 40.9510% | 100.00% | 946.9857 | 10.8 | 78,496,044 | 0.0640% | 99.98% | 77.9333 | 10.0 | 0.0196% | 99.97% | 323.3180 |
| 04 | 10.8 | 20,676,516 | 0.0524% | 98.40% | 5.2755 | 10.8 | 10,065,834 | 0.0126% | 81.80% | 1.7732 | 10.0 | 0.0001% | 76.27% | 1.7278 |
| 05 | 10.8 | 111,115,446 | 0.0926% | 100.00% | 61.0830 | 10.8 | 63,641,086 | 0.0092% | 99.85% | 8.8701 | 10.0 | 0.0001% | 56.29% | 0.9247 |
| 06 | 10.8 | 356,396,564 | 70.7456% | 100.00% | 3143.9166 | 10.8 | 78,571,822 | 0.2261% | 100.00% | 295.0117 | 10.0 | 0.0945% | 99.97% | 1469.0300 |
| 07 | 10.8 | 9,942,786 | 0.2915% | 99.64% | 10.7657 | 10.8 | 52,865,726 | 0.0101% | 98.97% | 6.5938 | 10.0 | 0.0002% | 94.25% | 3.8018 |
| 08 | 10.8 | 10,763,536 | 0.2497% | 99.54% | 8.5537 | 10.8 | 47,103,600 | 0.0067% | 91.82% | 2.7499 | 10.0 | 0.0008% | 66.42% | 2.2585 |
| 09 | 10.8 | 541,628,130 | 96.7444% | 100.00% | 3350.7269 | 10.8 | 21,166,616 | 5.9250% | 100.00% | 884.5224 | 10.0 | 4.7686% | 99.95% | 57.2028 |
| 10 | 10.8 | 5,668,752 | 0.7054% | 54.96% | 2.7434 | 10.8 | 4,089,390 | 0.1066% | 93.61% | 3.2721 | 10.0 | 0.0002% | 62.59% | 1.2704 |
| 11 | 10.8 | 11,176,696 | 0.2093% | 99.99% | 11.2174 | 10.8 | 41,913,790 | 0.0200% | 96.90% | 6.1212 | 10.0 | 0.0001% | 87.87% | 2.5211 |
| 12 | 10.8 | 16,151,064 | 27.1568% | 99.84% | 34.2213 | 10.8 | 40,226,610 | 0.0427% | 99.72% | 11.6552 | 10.0 | 0.0036% | 99.86% | 11.9888 |
| 13 | 10.8 | 14,881,342 | 0.2670% | 96.25% | 4.8453 | 10.8 | 7,335,290 | 0.0130% | 65.48% | 1.1660 | 10.0 | 0.0006% | 99.96% | 12.2455 |
| 14 | 10.8 | 13,884,580 | 3.0261% | 99.99% | 19.2685 | 10.8 | 21,795,080 | 0.1882% | 99.55% | 16.0357 | 10.0 | 0.0010% | 62.14% | 1.1220 |
| 15 | 10.8 | 12,387,588 | 65.9296% | 99.90% | 33.5811 | 10.8 | 3,105,472 | 0.3064% | 99.49% | 8.9043 | 10.0 | 0.0704% | 99.87% | 13.4546 |
| 16 | 10.8 | 13,025,918 | 9.2768% | 99.84% | 21.6711 | 10.8 | 19,495,766 | 0.3039% | 99.55% | 15.9261 | 10.0 | 0.0037% | 84.08% | 2.2149 |
| 17 | 10.8 | 16,384,556 | 13.2056% | 99.86% | 26.7111 | 10.8 | 32,164,850 | 0.1227% | 99.88% | 16.2674 | 10.0 | 0.0058% | 31.20% | 0.8448 |
| 18 | 2.7 | 112,794,866 | 19.1814% | 77.76% | 6.9234 | 2.7 | 35,616,834 | 0.0253% | 18.55% | 2.3147 | 10.0 | 0.0001% | 78.13% | 2.0642 |
| 19 | 2.7 | 93,872,216 | 4.1077% | 64.13% | 4.6651 | 2.7 | 35,775,154 | 0.0244% | 2.94% | 1.9927 | 10.0 | 0.0000% | 53.13% | 0.9280 |
| 20 | 2.7 | 82,493,546 | 0.0076% | 54.44% | 2.2178 | 2.7 | 36,822,898 | 0.0030% | 0.12% | 0.4650 | 10.0 | 0.0000% | 13.77% | 0.1814 |
| 21 | 2.7 | 92,874,254 | 11.0905% | 75.64% | 6.0904 | 2.7 | 22,813,548 | 0.0549% | 21.45% | 2.4093 | 10.0 | 0.0001% | 61.54% | 1.2370 |
| 22 | 2.7 | 106,391,154 | 0.2108% | 68.52% | 3.4031 | 2.7 | 37,797,862 | 0.0108% | 1.27% | 1.3720 | 10.0 | 0.0000% | 23.47% | 0.3527 |
| 23 | 2.7 | 135,954,444 | 0.0034% | 67.87% | 1.9198 | 2.7 | 36,422,158 | 0.0013% | 0.13% | 0.1155 | 10.0 | 0.0004% | 0.76% | 0.0201 |
| 24 | 2.7 | 68,556,412 | 0.7855% | 58.16% | 2.1196 | 2.7 | 32,646,446 | 0.0018% | 0.12% | 0.1614 | 10.0 | 0.0000% | 22.52% | 0.3292 |
| 25 | 2.7 | 201,614,222 | 47.7643% | 99.81% | 32.1785 | 2.7 | 34,297,778 | 0.0092% | 75.47% | 2.9363 | 10.0 | 0.0013% | 99.99% | 29.3693 |
| 26 | 2.7 | 642,140,604 | 92.8531% | 100.00% | 1621.9374 | 2.7 | 41,504,770 | 0.1649% | 99.89% | 124.7145 | 10.0 | 0.0694% | 99.99% | 1111.6309 |
| 27 | 2.7 | 28,519,004 | 0.0112% | 92.50% | 4.5196 | 2.7 | 34,802,762 | 0.0001% | 0.11% | 0.0351 | 10.0 | 0.0000% | 10.32% | 0.1326 |
| 28 | 2.7 | 9,792,364 | 0.1043% | 76.38% | 4.0190 | 2.7 | 22,201,210 | 0.0817% | 1.61% | 3.6903 | 10.0 | 0.0002% | 39.17% | 0.9250 |
| 29 | 2.7 | 175,344,692 | 75.5914% | 100.00% | 327.4946 | 2.7 | 33,195,792 | 0.0341% | 98.64% | 22.5668 | 10.0 | 0.0184% | 99.98% | 235.8581 |
| 30 | 2.7 | 25,325,652 | 0.0177% | 94.24% | 5.2230 | 2.7 | 36,375,652 | 0.0034% | 0.19% | 0.2462 | 10.0 | 0.0000% | 21.75% | 0.4325 |
| 31 | 2.7 | 51,558,410 | 0.2134% | 98.55% | 10.2587 | 2.7 | 48,321,376 | 0.0011% | 0.47% | 0.0996 | 10.0 | 0.0001% | 12.83% | 0.3015 |
| 32 | 2.7 | 1,638,196 | 17.5565% | 82.31% | 3.1014 | 2.7 | 2,855,228 | 2.6550% | 0.89% | 3.5194 | 10.0 | 0.0508% | 49.21% | 7.7024 |
| 34 | 5.4 | 146,373,482 | 90.6535% | 100.00% | 7359.4100 | 5.4 | 10,605,224 | 0.7548% | 100.00% | 176.2427 | 10.0 | 0.1662% | 99.98% | 842.7465 |
| 35 | 5.4 | 749,392 | 13.5239% | 33.49% | 1.1914 | 5.4 | 4,854,666 | 0.0509% | 4.67% | 1.2298 | 10.0 | 0.0041% | 32.24% | 0.5876 |
| 36 | 5.4 | 66,509,586 | 0.1866% | 99.96% | 23.6832 | 5.4 | 111,047,572 | 0.0001% | 7.79% | 0.1038 | 10.0 | 0.0003% | 57.51% | 1.0422 |
| 37 | 5.4 | 9,834,452 | 60.1041% | 98.86% | 10.6667 | 5.4 | 4,406,210 | 0.0336% | 74.81% | 2.0089 | 10.0 | 0.0201% | 56.10% | 1.2234 |
| 38 | 5.4 | 43,022,752 | 97.0263% | 100.00% | 252.6044 | 5.4 | 1,123,198 | 0.9702% | 99.99% | 23.4852 | 10.0 | 0.5773% | 99.85% | 17.2460 |
| 39 | 5.4 | 1,340,114 | 2.0089% | 4.79% | 0.4514 | 5.4 | 3,976,390 | 0.0153% | 2.83% | 0.3394 | 10.0 | 0.0009% | 24.95% | 0.4494 |
| 40 | 5.4 | 3,628,686 | 5.6932% | 24.91% | 0.8514 | 5.4 | 4,912,264 | 0.0177% | 9.88% | 0.5260 | 10.0 | 0.0008% | 19.53% | 0.3346 |
| 41 | 5.4 | 29,918,078 | 83.4096% | 99.86% | 48.3122 | 5.4 | 5,950,044 | 0.1372% | 98.88% | 10.4956 | 10.0 | 0.1244% | 99.91% | 25.1505 |
| 42 | 5.4 | 63,305,808 | 92.6539% | 99.99% | 242.1007 | 5.4 | 2,034,818 | 0.4434% | 99.61% | 17.8696 | 10.0 | 0.1145% | 99.97% | 75.3447 |
| 43 | 5.4 | 28,502,096 | 2.0792% | 98.90% | 8.6775 | 5.4 | 21,291,192 | 0.0006% | 20.84% | 0.2769 | 10.0 | 0.0008% | 43.21% | 0.8251 |
| 44 | 5.4 | 23,678,350 | 0.1632% | 73.65% | 2.0966 | 5.4 | 15,802,662 | 0.0008% | 13.19% | 0.1653 | 10.0 | 0.0004% | 31.84% | 0.5936 |
| 45 | 5.4 | 28,763,804 | 3.0793% | 99.93% | 15.9794 | 5.4 | 19,682,236 | 0.0196% | 96.32% | 4.9701 | 10.0 | 0.0012% | 97.70% | 5.9600 |
| 46 | 5.4 | 48,033,536 | 10.5088% | 100.00% | 158.1837 | 5.4 | 31,002,106 | 0.0034% | 81.51% | 2.4673 | 10.0 | 0.0041% | 99.96% | 41.5512 |
| 47 | 5.4 | 32,374,174 | 6.1377% | 99.89% | 17.6006 | 5.4 | 20,075,024 | 0.0017% | 42.98% | 0.7470 | 10.0 | 0.0014% | 84.73% | 2.6273 |
| 48 | 5.4 | 33,545,522 | 0.8612% | 95.79% | 6.1428 | 5.4 | 24,579,728 | 0.0003% | 8.20% | 0.1238 | 10.0 | 0.0008% | 54.43% | 1.1477 |
| 49 | 5.4 | 41,228,324 | 1.0147% | 99.88% | 18.1782 | 5.4 | 29,981,390 | 0.0003% | 11.99% | 0.1741 | 10.0 | 0.0005% | 91.95% | 3.3415 |
| 50 | 5.4 | 12,177,692 | 71.4756% | 99.90% | 121.7697 | 5.4 | 22,754,546 | 0.0264% | 99.44% | 11.6945 | 10.0 | 0.0421% | 99.96% | 51.3386 |
| 51 | 5.4 | 6,105,636 | 46.9134% | 99.78% | 28.5245 | 5.4 | 21,784,580 | 0.0109% | 90.56% | 3.7560 | 10.0 | 0.0058% | 99.95% | 25.2443 |
| 52 | 5.4 | 3,830,746 | 4.8117% | 90.86% | 4.5674 | 5.4 | 24,464,202 | 0.0008% | 14.20% | 0.2901 | 10.0 | 0.0005% | 89.58% | 2.9335 |
| 53 | 5.4 | 6,556,106 | 32.5097% | 100.00% | 72.9775 | 5.4 | 25,666,594 | 0.0055% | 82.54% | 2.7256 | 10.0 | 0.0033% | 99.96% | 47.4457 |

| | | | | | | | | | | | | | | |
|---|---|---|---|---|---|---|---|---|---|---|---|---|---|---|
| 54 | 5.4 | 45,398,712 | 88.7216% | 100.00% | 1074.7572 | 5.4 | 28,016,362 | 0.0849% | 100.00% | 52.9310 | 10.0 | 0.0896% | 100.00% | 843.4279 |
| 55 | 5.4 | 103,807,862 | 94.1763% | 100.00% | 3139.9488 | 5.4 | 25,661,784 | 0.2476% | 100.00% | 139.5156 | 10.0 | 0.1555% | 99.99% | 1233.6985 |
| 56 | 5.4 | 5,072,904 | 21.2207% | 99.86% | 38.2663 | 5.4 | 25,257,896 | 0.0029% | 59.39% | 1.3265 | 10.0 | 0.0020% | 99.96% | 37.2734 |
| 57 | 5.4 | 26,651,568 | 81.5262% | 100.00% | 764.1384 | 5.4 | 23,771,526 | 0.0475% | 99.90% | 25.3226 | 10.0 | 0.0392% | 100.00% | 773.4338 |
| 58 | 5.4 | 119,250,340 | 89.4189% | 100.00% | 1499.0878 | 5.4 | 21,737,454 | 1.3970% | 99.99% | 481.7897 | 10.0 | 0.8035% | 99.96% | 18.0897 |
| 59 | 5.4 | 13,876,992 | 73.0525% | 99.98% | 94.7249 | 5.4 | 6,897,404 | 0.2907% | 99.98% | 23.3441 | 10.0 | 0.0756% | 60.08% | 1.0516 |
| 60 | 5.4 | 5,064,692 | 0.7576% | 81.74% | 2.8944 | 5.4 | 28,676,654 | 0.0022% | 7.97% | 0.4627 | 10.0 | 0.0004% | 46.13% | 0.9281 |
| 61 | 5.4 | 3,237,272 | 2.4584% | 94.59% | 4.1871 | 5.4 | 12,599,798 | 0.0007% | 14.80% | 0.1898 | 10.0 | 0.0003% | 38.29% | 0.6626 |
| 62 | 5.4 | 6,685,592 | 12.8404% | 99.96% | 46.9370 | 5.4 | 18,066,044 | 0.0082% | 76.60% | 2.4364 | 10.0 | 0.0048% | 99.89% | 11.2049 |
| 63 | 5.4 | 6,482,172 | 1.7973% | 96.84% | 5.4459 | 5.4 | 23,768,308 | 0.0006% | 21.74% | 0.2900 | 10.0 | 0.0005% | 77.05% | 2.1138 |
| 64 | 5.4 | 9,466,060 | 0.2669% | 96.99% | 4.5785 | 5.4 | 22,213,606 | 0.0001% | 3.55% | 0.0437 | 10.0 | 0.0001% | 49.20% | 0.8110 |
| 65 | 5.4 | 5,783,354 | 0.3323% | 76.84% | 1.9841 | 5.4 | 24,375,006 | 0.0001% | 2.16% | 0.0297 | 10.0 | 0.0000% | 47.11% | 0.6932 |
| 66 | 5.4 | 5,970,510 | 29.3245% | 99.91% | 30.3217 | 5.4 | 22,078,330 | 0.0951% | 99.92% | 16.9899 | 10.0 | 0.0290% | 92.86% | 4.1057 |
| 67 | 5.4 | 8,697,652 | 47.3466% | 100.00% | 768.4558 | 5.4 | 23,501,026 | 0.1764% | 100.00% | 88.7774 | 10.0 | 0.1034% | 99.97% | 490.0341 |
| 68 | 5.4 | 25,258,898 | 76.1765% | 100.00% | 4894.2700 | 5.4 | 28,494,342 | 0.8557% | 100.00% | 475.1583 | 10.0 | 0.5174% | 99.96% | 177.1438 |
| 69 | 5.4 | 141,070,280 | 96.2193% | 100.00% | 7846.8553 | 5.4 | 28,135,628 | 13.5652% | 100.00% | 4153.0717 | 10.0 | 8.8958% | 100.00% | 6361.1653 |
| 70 | 5.4 | 5,686,808 | 0.0154% | 77.96% | 1.8157 | 5.4 | 24,052,452 | 0.0000% | 0.54% | 0.0102 | 10.0 | 0.0010% | 99.68% | 9.4708 |
| 71 | 5.4 | 9,016,134 | 18.4892% | 100.00% | 506.1078 | 5.4 | 30,946,146 | 0.0458% | 99.95% | 30.1117 | 10.0 | 0.0199% | 99.96% | 26.0630 |
| 72 | 5.4 | 7,900,462 | 7.0146% | 99.99% | 23.4415 | 5.4 | 10,547,490 | 0.0016% | 25.25% | 0.3908 | 10.0 | 0.0018% | 99.96% | 18.4789 |
| 73 | 5.4 | 8,211,094 | 39.8226% | 99.99% | 87.6919 | 5.4 | 20,473,706 | 0.1506% | 99.99% | 38.6144 | 10.0 | 0.0774% | 99.96% | 16.5545 |
| 74 | 5.4 | 13,940,912 | 1.6234% | 54.47% | 1.7018 | 5.4 | 10,982,782 | 0.0001% | 1.71% | 0.0198 | 10.0 | 0.0023% | 99.95% | 43.3575 |
| 75 | 5.4 | 53,210,982 | 59.1457% | 100.00% | 148.0972 | 5.4 | 12,675,448 | 0.0103% | 86.27% | 2.8130 | 10.0 | 0.0057% | 99.62% | 8.0070 |
| 76 | 5.4 | 5,350,422 | 4.1008% | 38.45% | 0.6652 | 5.4 | 4,988,872 | 0.0003% | 2.41% | 0.0256 | 10.0 | 0.0005% | 99.45% | 10.3293 |
| 77 | 5.4 | 80,704,356 | 27.2584% | 99.68% | 20.6439 | 5.4 | 41,708,526 | 0.0016% | 67.15% | 1.3075 | 10.0 | 0.0012% | 99.59% | 10.0565 |
| 78 | 5.4 | 87,431,520 | 63.3773% | 99.92% | 55.5415 | 5.4 | 21,631,534 | 0.0194% | 93.61% | 4.5826 | 10.0 | 0.0063% | 89.19% | 2.5408 |
| 79 | 5.4 | 27,533,572 | 0.3070% | 45.27% | 0.7658 | 5.4 | 16,031,304 | 0.0000% | 0.12% | 0.0042 | 10.0 | 0.0003% | 98.62% | 6.1102 |
| 80 | 5.4 | 3,872,932 | 72.5768% | 99.34% | 14.4106 | 5.4 | 1,250,890 | 0.0831% | 63.00% | 1.5866 | 10.0 | 0.1389% | 97.92% | 5.2081 |
| 81 | 5.4 | 4,648,190 | 7.6058% | 98.09% | 9.4652 | 5.4 | 3,511,362 | 0.0003% | 2.11% | 0.0210 | 10.0 | 0.0007% | 99.94% | 13.4037 |

**Protocol Optimization**

| | | | | | | | | | | | |
|---|---|---|---|---|---|---|---|---|---|---|---|
| 02 | | | | | | 2.7 | 16,408,970 | 0.1012% | 88.32% | 4.8521 | |
| 03 | | | | | | 2.7 | 47,083,344 | 0.0520% | 99.96% | 40.6153 | |
| 04 | | | | | | 2.7 | 57,543,194 | 0.0027% | 44.43% | 0.7633 | |
| 05 | | | | | | 2.7 | 59,021,696 | 0.0019% | 41.02% | 0.6959 | |
| 06 | | | | | | 2.7 | 43,005,642 | 0.1520% | 100.00% | 114.7798 | |
| 07 | 2.7 | 6,807,324 | 0.2123% | 92.42% | 3.2703 | 2.7 | 59,245,398 | 0.0048% | 93.44% | 4.4251 | |
| 08 | 2.7 | 7,133,858 | 0.2009% | 95.80% | 4.2571 | 2.7 | 53,691,494 | 0.0057% | 91.36% | 4.0402 | |
| 09 | 2.7 | 534,740,522 | 97.9777% | 100.00% | 2185.2300 | 2.7 | 17,555,514 | 6.0440% | 100.00% | 378.9593 | |
| 10 | | | | | | 2.7 | 17,409,862 | 0.1325% | 84.16% | 5.2398 | Input volume: 2.7 µl vs 10.8 µl |
| 11 | 2.7 | 58 | 0.0000% | 0.00% | 0.0000 | 2.7 | 1,618 | 0.0618% | 0.10% | 0.0010 | |
| 12 | 2.7 | 17,074,722 | 30.2408% | 99.61% | 15.5067 | 2.7 | 47,310,582 | 0.0206% | 98.08% | 9.1497 | |
| 13 | | | | | | 2.7 | 48,064,580 | 0.0035% | 41.59% | 0.7463 | |
| 14 | 2.7 | 12,514,502 | 4.9961% | 98.33% | 7.4607 | 2.7 | 28,491,880 | 0.1188% | 99.49% | 18.5844 | |
| 15 | | | | | | 2.7 | 18,079,484 | 0.2210% | 98.69% | 10.7718 | |
| 16 | 2.7 | 8,061,366 | 18.2108% | 94.00% | 4.9191 | 2.7 | 15,778,084 | 0.1176% | 98.85% | 12.2731 | |
| 17 | 2.7 | 17,317,738 | 12.2331% | 99.49% | 11.8779 | 2.7 | 45,806,156 | 0.0700% | 99.44% | 17.8944 | |
| 50 | 2.7 | 5,302,774 | 71.9360% | 99.74% | 56.2274 | 2.7 | 9,440,676 | 0.0260% | 94.59% | 4.86135 | |
| | | 6,874,918 | 71.0528% | 99.90% | 68.0711 | | 13,313,870 | 0.0267% | 97.44% | 6.94325 | |
| 51 | 2.7 | 2,651,528 | 45.9457% | 99.27% | 12.73 | 2.7 | 9,956,018 | 0.0106% | 66.31% | 1.68468 | |
| | | 3,454,108 | 47.6123% | 99.75% | 16.0504 | | 11,828,562 | 0.0112% | 75.74% | 2.11945 | |
| 52 | 2.7 | 1,812,456 | 5.1702% | 78.61% | 2.70916 | 2.7 | 11,905,088 | 0.0011% | 7.45% | 0.166271 | |
| | | 2,018,290 | 4.4788% | 68.52% | 1.99716 | | 12,559,114 | 0.0006% | 8.10% | 0.129552 | |
| 53 | 2.7 | 2,661,912 | 32.8762% | 99.97% | 33.6088 | 2.7 | 11,393,230 | 0.0058% | 56.44% | 1.28014 | |
| | | 3,894,194 | 32.2177% | 99.91% | 40.2593 | | 14,273,364 | 0.0054% | 64.91% | 1.47253 | Input volume: 2.7 µl vs 5.4 µl |

| | | | | | | | | | | | |
|---|---|---|---|---|---|---|---|---|---|---|---|
| 54 | 2.7 | 16,862,842 | 88.9164% | 100.00% | 554.182 | 2.7 | 10,453,800 | 0.0886% | 99.98% | 21.1365 | Input volume: 2.7 µl vs 5.4 µl |
| | | 28,535,870 | 88.4747% | 99.98% | 623.171 | | 17,562,562 | 0.0827% | 100.00% | 32.3643 | |
| 55 | 2.7 | 50,308,172 | 94.4399% | 100.00% | 1763.58 | 2.7 | 12,242,678 | 0.2470% | 99.97% | 68.163 | |
| | | 53,499,690 | 93.7940% | 100.00% | 1910.21 | | 13,419,106 | 0.2482% | 100.00% | 75.0258 | |
| 56 | 2.7 | 2,407,594 | 21.5466% | 99.75% | 18.4627 | 2.7 | 11,510,576 | 0.0029% | 34.62% | 0.621309 | |
| | | 2,665,310 | 20.8997% | 99.82% | 20.063 | | 13,747,320 | 0.0029% | 39.51% | 0.716082 | |
| 57 | 2.7 | 12,058,338 | 80.8714% | 100.00% | 375.207 | 2.7 | 10,977,398 | 0.0477% | 99.15% | 11.8002 | |
| | | 14,593,230 | 81.9503% | 100.00% | 438.47 | | 12,794,128 | 0.0474% | 99.35% | 13.7123 | |
| 42 | 5.4 | 63,305,808 | 92.6539% | 99.99% | 242.1007 | 5.4 | 2,034,818 | 0.4434% | 99.61% | 17.8696 | With post-added carrier RNA |
| 43 | 5.4 | 28,502,096 | 2.0792% | 98.90% | 8.6775 | 5.4 | 21,291,192 | 0.0006% | 20.84% | 0.2769 | |
| 44 | 5.4 | 23,678,350 | 0.1632% | 73.65% | 2.0966 | 5.4 | 15,802,662 | 0.0008% | 13.19% | 0.1653 | |
| 45 | 5.4 | 28,763,804 | 3.0793% | 99.93% | 15.9794 | 5.4 | 19,682,236 | 0.0196% | 96.32% | 4.9701 | |
| 46 | 5.4 | 48,033,536 | 10.5088% | 100.00% | 158.1837 | 5.4 | 31,002,106 | 0.0034% | 81.51% | 2.4673 | |
| 47 | 5.4 | 32,374,174 | 6.1377% | 99.89% | 17.6006 | 5.4 | 20,075,024 | 0.0017% | 42.98% | 0.747 | |
| 48 | 5.4 | 33,545,522 | 0.8612% | 95.79% | 6.1428 | 5.4 | 24,579,728 | 0.0003% | 8.20% | 0.1238 | |
| 49 | 5.4 | 41,228,324 | 1.0147% | 99.88% | 18.1782 | 5.4 | 29,981,390 | 0.0003% | 11.99% | 0.1741 | |
| Y1 | 5.4 | 1,267,440 | 56.3684% | 93.93% | 5.5805 | 5.4 | 751,372 | 0.0527% | 25.64% | 0.5249 | RNA extraction with carrier RNA |
| Y2 | 5.4 | 75,641,774 | 86.5550% | 100.00% | 4395.5348 | 5.4 | 5,811,906 | 0.4477% | 99.98% | 58.3571 | |
| Y3 | 5.4 | 40,132,820 | 38.8485% | 100.00% | 145.8907 | 5.4 | 29,519,046 | 0.0144% | 98.34% | 9.2615 | |
| Y4 | 5.4 | 21,566,432 | 11.2625% | 99.85% | 32.3432 | 5.4 | 22,669,654 | 0.0036% | 70.52% | 1.6935 | |
| Y5 | 5.4 | 9,021,160 | 2.7007% | 86.56% | 3.0329 | 5.4 | 8,699,170 | 0.0011% | 7.99% | 0.1337 | |
| Y6 | 5.4 | 18,074,214 | 97.7771% | 100.00% | 968.249 | 5.4 | 18,398 | 30.6175% | 99.58% | 13.0292 | |

#### SHERRY Metagenomic Analysis

| Sample ID | Total Reads | Human Ratio | Nonhuman Reads | Viral Ratio (to Nonhuman) | Fungal Ratio (to Nonhuman) | Bacterial Ratio (to Nonhuman) |
|---|---|---|---|---|---|---|
| 002 | 2,468,936 | 18.9023% | 494,070 | 1.1173% | 4.5481% | 52.3414% |
| 003 | 39,092,827 | 0.3152% | 8,023,735 | 0.5856% | 1.8087% | 70.8749% |
| 004 | 5,013,516 | 0.2248% | 787,922 | 0.3187% | 1.8070% | 75.9708% |
| 005 | 31,683,662 | 0.3376% | 2,361,502 | 0.4108% | 3.6783% | 80.0814% |
| 006 | 39,139,616 | 0.1557% | 4,297,298 | 2.4188% | 2.3355% | 68.3550% |
| 007 | 26,331,498 | 0.2152% | 3,490,992 | 0.2330% | 2.1136% | 80.4029% |
| 008 | 23,484,173 | 0.2497% | 4,366,334 | 0.3063% | 2.2150% | 81.7824% |
| 009 | 10,092,707 | 23.5251% | 1,513,285 | 41.7013% | 6.3245% | 16.8219% |
| 010 | 1,979,521 | 10.0166% | 292,498 | 1.0496% | 5.0438% | 34.2553% |
| 011 | 20,857,619 | 0.5797% | 4,936,618 | 0.2005% | 1.8073% | 69.1540% |
| 012 | 19,935,080 | 5.6740% | 2,872,201 | 0.4993% | 2.5371% | 68.0528% |
| 013 | 3,655,972 | 0.2099% | 995,977 | 0.3084% | 1.4819% | 84.9455% |
| 014 | 10,463,234 | 7.4030% | 1,012,355 | 1.4751% | 10.4525% | 30.0168% |
| 015 | 1,487,902 | 13.7522% | 132,205 | 2.9220% | 13.6674% | 29.6290% |
| 016 | 9,206,968 | 12.8453% | 919,591 | 1.3036% | 15.7630% | 30.0165% |
| 017 | 15,710,532 | 7.5051% | 3,373,926 | 0.4682% | 51.9567% | 12.7691% |
| 018 | 17,808,417 | 0.4963% | 273,574 | 0.2409% | 30.0961% | 38.0548% |
| 019 | 17,887,577 | 0.2036% | 1,126,424 | 0.1544% | 9.2432% | 74.0607% |
| 020 | 18,411,449 | 0.2011% | 226,583 | 0.1289% | 27.5643% | 42.5058% |
| 021 | 11,406,774 | 2.1513% | 384,049 | 0.8835% | 22.7106% | 43.4989% |
| 022 | 18,898,931 | 0.1262% | 467,215 | 0.1988% | 17.8305% | 52.5204% |
| 023 | 18,211,079 | 0.0968% | 2,612,059 | 0.2827% | 8.7084% | 61.0145% |
| 024 | 16,323,223 | 0.3676% | 14,482,545 | 98.9738% | 0.4180% | 0.2953% |
| 025 | 17,148,889 | 0.2921% | 3,314,711 | 0.2424% | 7.8022% | 61.3573% |
| 026 | 20,752,385 | 0.4237% | 1,924,866 | 2.1696% | 10.4624% | 69.1715% |
| 027 | 17,401,381 | 0.1131% | 309,059 | 0.5015% | 28.3328% | 50.8369% |
| 028 | 11,100,605 | 86.2836% | 1,187,883 | 0.7710% | 31.2298% | 17.5335% |
| 029 | 16,597,896 | 0.7660% | 1,480,193 | 0.6239% | 12.0163% | 59.8970% |
| 030 | 18,187,826 | 31.8460% | 1,834,920 | 0.4200% | 24.2711% | 38.0481% |
| 031 | 24,160,688 | 0.4161% | 1,035,181 | 0.8847% | 17.2940% | 43.7442% |
| 032 | 1,427,614 | 66.1264% | 338,880 | 1.1101% | 38.9633% | 21.9154% |
| 034 | 5,300,922 | 1.6809% | 835,001 | 5.0602% | 2.7047% | 69.4854% |
| 035 | 2,426,574 | 78.7935% | 619,841 | 2.0623% | 7.5047% | 11.5150% |
| 036 | 55,506,497 | 0.1535% | 5,513,710 | 0.3863% | 4.1224% | 74.7251% |
| 037 | 2,202,567 | 21.9957% | 184,865 | 0.8087% | 19.5094% | 48.3575% |
| 038 | 561,441 | 26.0678% | 37,991 | 14.4455% | 14.4166% | 35.2662% |
| 039 | 1,987,593 | 41.1263% | 263,095 | 1.0650% | 9.6840% | 36.3526% |
| 040 | 2,455,378 | 33.7620% | 286,824 | 1.1467% | 9.7931% | 36.8463% |
| 041 | 2,974,091 | 15.9757% | 263,758 | 2.2672% | 7.9171% | 47.9015% |
| 042 | 1,017,101 | 52.3014% | 71,365 | 6.9740% | 9.9110% | 23.9879% |
| 043 | 10,642,295 | 0.2241% | 1,419,615 | 0.1659% | 3.4087% | 64.9659% |
| 044 | 7,898,876 | 0.5322% | 1,579,626 | 0.1700% | 2.7013% | 65.0570% |
| 045 | 9,838,150 | 0.2754% | 1,952,079 | 0.2996% | 2.6873% | 65.3771% |
| 046 | 15,496,315 | 0.0703% | 2,672,772 | 0.2713% | 2.4649% | 72.5376% |
| 047 | 10,034,335 | 0.4773% | 1,278,475 | 0.1494% | 3.2776% | 65.1730% |
| 048 | 12,286,185 | 0.1687% | 2,318,133 | 0.3839% | 2.4469% | 70.7022% |
| 049 | 14,986,090 | 0.2644% | 2,696,632 | 0.1065% | 2.3671% | 64.6335% |
| 050 | 16,097,611 | 0.7683% | 3,220,837 | 0.3401% | 2.9477% | 69.9487% |
| 051 | 15,870,299 | 1.6305% | 4,544,027 | 0.5113% | 2.1790% | 89.7928% |
| 052 | 18,511,658 | 1.4084% | 3,064,879 | 0.2117% | 3.1124% | 64.0877% |
| 053 | 18,529,912 | 0.5910% | 2,732,709 | 0.1599% | 3.0647% | 60.0824% |
| 054 | 19,235,081 | 0.2929% | 3,265,342 | 0.6110% | 2.6649% | 60.6792% |
| 055 | 18,952,231 | 0.1798% | 3,837,017 | 1.4074% | 2.4665% | 63.5462% |
| 056 | 18,384,236 | 0.1544% | 3,581,067 | 0.0865% | 2.2282% | 66.5599% |
| 057 | 17,374,462 | 0.3177% | 3,346,679 | 0.4133% | 2.4326% | 67.0528% |
| 058 | 10,868,727 | 2.2796% | 638,484 | 23.6040% | 7.8437% | 40.3385% |
| 059 | 3,448,702 | 5.7016% | 270,289 | 3.3819% | 6.0857% | 28.4902% |
| 060 | 14,338,327 | 1.7012% | 2,339,720 | 0.1854% | 2.5786% | 59.6549% |
| 061 | 6,299,899 | 0.0922% | 939,891 | 0.1568% | 4.3892% | 63.9443% |
| 062 | 9,033,022 | 0.5409% | 1,399,198 | 0.2913% | 3.3986% | 71.5118% |
| 063 | 11,884,154 | 0.1248% | 1,141,437 | 0.1111% | 5.6198% | 62.0442% |
| 064 | 11,106,803 | 0.1482% | 1,121,648 | 0.1078% | 3.6808% | 58.2269% |
| 065 | 12,187,503 | 0.1927% | 1,167,864 | 0.4349% | 4.2675% | 79.3624% |
| 066 | 11,039,165 | 7.9590% | 2,729,856 | 1.0843% | 2.2071% | 69.3473% |
| 067 | 11,750,513 | 0.2689% | 1,612,208 | 1.4605% | 4.0225% | 62.7300% |
| 068 | 14,247,171 | 0.8420% | 2,602,295 | 4.7992% | 2.5559% | 54.0959% |
| 069 | 14,067,814 | 8.9599% | 3,831,180 | 49.3960% | 1.4864% | 31.2107% |
| 070 | 12,026,226 | 0.1070% | 1,187,881 | 0.2206% | 4.4802% | 62.1201% |
| 071 | 15,473,073 | 6.8376% | 2,455,028 | 0.4899% | 2.9558% | 73.1784% |
| 072 | 5,273,745 | 0.4997% | 1,036,507 | 0.1826% | 3.2783% | 67.0467% |
| 073 | 10,236,853 | 3.2561% | 1,503,578 | 1.2516% | 5.0292% | 68.1341% |
| 074 | 5,491,391 | 0.1041% | 1,496,732 | 0.1725% | 2.5978% | 81.1353% |
| 075 | 6,337,724 | 0.0682% | 849,815 | 0.2279% | 2.8418% | 67.1447% |
| 076 | 2,494,436 | 0.1104% | 104,031 | 0.1278% | 10.5382% | 78.3901% |
| 077 | 20,854,263 | 0.1378% | 664,662 | 0.0927% | 7.6522% | 69.5597% |
| 078 | 10,815,767 | 1.2924% | 1,170,985 | 0.6318% | 7.6884% | 69.1229% |
| 079 | 8,015,652 | 0.5025% | 1,076,912 | 0.2192% | 4.3354% | 68.8388% |
| 080 | 625,445 | 49.7480% | 65,519 | 1.8468% | 16.5998% | 31.2825% |
| 081 | 1,755,681 | 0.0239% | 107,785 | 0.0130% | 7.8610% | 76.4392% |
| Y1 | 1,710,655 | 24.5093% | 73,250 | 0.3741% | 3.6642% | 30.6198% |
| Y2 | 2,905,953 | 2.2314% | 205,752 | 7.0434% | 4.0476% | 50.6624% |
| Y3 | 14,759,523 | 1.4268% | 736,811 | 0.4001% | 8.2238% | 54.3242% |
| Y4 | 11,334,827 | 3.9014% | 989,139 | 0.1116% | 5.1537% | 54.7062% |
| Y5 | 4,349,585 | 4.7464% | 811,926 | 0.1020% | 3.4745% | 59.2098% |
| Y6 | 884,080 | 49.9126% | 4,143 | 63.9150% | 2.0517% | 9.5342% |
| 083 | 3,941,854 | 0.2700% | 424,059 | 0.3443% | 3.2672% | 61.3620% |
| 085 | 18,024,622 | 0.2132% | 1,683,305 | 0.1428% | 6.7048% | 74.7914% |
| 086 | 22,841,425 | 0.1809% | 1,998,054 | 0.4592% | 4.1101% | 74.0862% |
| 088 | 16,051,454 | 0.1610% | 2,370,305 | 0.4237% | 3.3140% | 71.4821% |
| 089 | 90,048,297 | 12.5490% | 6,824,623 | 0.2910% | 8.3339% | 61.2114% |
| 093 | 48,439,555 | 0.1485% | 11,315,945 | 0.2669% | 1.9454% | 78.7332% |
| 094 | 29,001,384 | 3.1525% | 1,801,276 | 0.2588% | 8.4963% | 57.6490% |
| 095 | 36,614,283 | 0.2029% | 9,966,114 | 0.2932% | 2.5249% | 78.9526% |
| 096 | 17,693,004 | 0.3554% | 2,728,927 | 0.1660% | 2.3488% | 64.3970% |

| | | | | | | |
|---|---|---|---|---|---|---|
| 102 | 38,845,063 | 1.2532% | 7,315,003 | 0.2669% | 5.1053% | 62.5628% |
| 113 | 28,962,515 | 26.7526% | 1,265,738 | 0.6692% | 12.9786% | 41.2912% |
| 121 | 61,048,022 | 0.4617% | 9,471,265 | 0.0885% | 2.7632% | 56.9690% |
| 122 | 62,531,557 | 0.1011% | 3,642,880 | 0.1839% | 3.6620% | 71.8376% |
| 123 | 18,038,879 | 0.1338% | 727,333 | 0.1730% | 3.7015% | 57.0275% |
| 125 | 147,956 | 45.0101% | 70,914 | 1.3213% | 4.1092% | 35.8660% |
| 128 | 23,954,438 | 4.5390% | 2,512,033 | 0.8108% | 3.9145% | 66.6291% |
| 145 | 3,694,592 | 21.1780% | 837,066 | 16.0463% | 9.7836% | 18.0955% |
| 146 | 1,909,718 | 6.3736% | 130,696 | 0.3688% | 8.8924% | 74.2624% |
| 147 | 3,497,422 | 15.3439% | 271,297 | 0.5175% | 5.3672% | 51.9342% |
| 148 | 65,982,369 | 3.8598% | 10,604,568 | 0.2198% | 3.1893% | 68.3702% |
| 149 | 59,348,449 | 13.5240% | 4,498,775 | 0.6450% | 4.3098% | 49.6394% |
| 150 | 49,972,286 | 16.9918% | 3,665,622 | 0.6884% | 7.2268% | 50.4646% |
| 154 | 1,100,444 | 24.0664% | 221,335 | 6.1987% | 6.3406% | 22.3616% |
| 167 | 36,444,346 | 0.1215% | 9,913,749 | 0.2558% | 2.7010% | 78.5594% |
| 168 | 3,315,967 | 0.1420% | 1,155,851 | 0.2136% | 4.8643% | 72.8685% |
| 169 | 657,575 | 6.0191% | 107,793 | 1.2366% | 7.6406% | 34.2304% |
| 170 | 47,197,271 | 0.1587% | 6,999,442 | 0.1974% | 7.5890% | 74.0438% |
| 174 | 11,735,414 | 19.1732% | 2,646,320 | 0.5561% | 59.8146% | 18.5157% |
| 177 | 4,080,078 | 50.2751% | 343,623 | 2.4652% | 10.8156% | 33.2268% |
| 180 | 28,253,580 | 0.2115% | 1,001,601 | 0.0532% | 11.7777% | 60.1394% |
| 181 | 8,753,533 | 3.3779% | 578,560 | 0.2987% | 12.2637% | 58.6088% |
| 182 | 14,423,027 | 1.2391% | 624,906 | 0.2301% | 19.8494% | 45.3302% |
| 183 | 31,024,516 | 0.6847% | 2,486,586 | 0.1181% | 5.4932% | 59.8985% |
| 184 | 28,167,697 | 0.0957% | 6,152,701 | 0.1216% | 2.6230% | 54.4071% |
| 185 | 32,399,243 | 0.1434% | 10,373,292 | 0.1358% | 2.0667% | 79.5981% |
| 186 | 2,690,386 | 14.6166% | 343,916 | 0.6682% | 6.2704% | 47.0010% |
| 189 | 55,864,509 | 0.2723% | 1,873,722 | 0.7107% | 6.2748% | 72.9392% |
| 192 | 47,469,542 | 0.5586% | 1,901,991 | 0.2362% | 5.5677% | 73.5656% |
| 193 | 59,688,959 | 0.3591% | 13,047,108 | 0.1418% | 2.6803% | 58.3770% |
| 194 | 37,441,392 | 0.3134% | 15,221,187 | 0.2546% | 1.5260% | 74.6436% |
| 195 | 39,710,104 | 0.3632% | 1,308,850 | 0.9599% | 4.9410% | 60.9372% |
| 202 | 44,813,797 | 1.0700% | 2,811,087 | 0.0547% | 4.6688% | 36.4013% |
| 212 | 19,176,613 | 2.9842% | 1,346,975 | 0.2861% | 3.5996% | 43.8362% |
| 215 | 3,288,098 | 61.9402% | 787,932 | 0.5943% | 3.5762% | 23.5500% |
| 217 | 50,415,096 | 0.5448% | 4,264,261 | 0.0804% | 4.6439% | 50.4234% |
| 230 | 64,725,702 | 0.1840% | 4,858,865 | 4.0872% | 2.8553% | 57.3845% |
| 234 | 1,293,834 | 58.3609% | 181,154 | 0.3930% | 2.6099% | 23.9603% |
| 235 | 392,949 | 24.5952% | 19,768 | 0.2529% | 2.0285% | 22.2177% |
| 241 | 3,851,504 | 18.0306% | 1,320,663 | 0.3024% | 2.9584% | 32.6097% |
| 302 | 27,406,423 | 10.9655% | 3,954,842 | 0.1874% | 4.1846% | 33.8902% |
| 303 | 72,088,145 | 0.2610% | 13,874,159 | 0.1691% | 2.0673% | 68.8517% |

| MINERVA Supplementary Sample | | | | |
|---|---|---|---|---|
| Sample ID | Total Reads | Mapping Ratio | Dedup Coverage | Dedup Depth |
| 122 | 39,411,066 | 8.5232% | 99.39% | 10.5703 |
| 123 | 36,100,522 | 74.2424% | 100.00% | 213.5296 |
| 148 | 139,565,182 | 46.5267% | 95.02% | 14.5469 |
| 154 | 252,038,586 | 98.9726% | 100.00% | 420.7170 |
| 189 | 374,145,632 | 93.9090% | 100.00% | 1,582.2800 |
| 192 | 60,506,220 | 12.5611% | 98.28% | 11.9391 |
| 193 | 72,845,794 | 13.1650% | 99.83% | 24.8392 |
| 194 | 113,929,174 | 3.5088% | 94.84% | 6.6863 |
| 195 | 231,325,932 | 55.7130% | 99.93% | 128.3160 |
| 202 | 95,943,060 | 1.7287% | 99.32% | 17.7605 |
| 247 | 133,758,300 | 9.0448% | 98.10% | 12.7086 |
| Y14 | 175,555,840 | 80.2967% | 84.94% | 25.7050 |
| Y23 | 99,919,844 | 0.3094% | 27.33% | 2.9570 |
| Y31 | 50,920,168 | 75.4457% | 95.71% | 27.4759 |
| Y32 | 98,258,964 | 0.4034% | 84.61% | 6.1846 |
| Y37 | 30,814,630 | 10.8579% | 55.43% | 2.1652 |

Supplementary Appendix - Protocol

### Protocol: MINERVA – a rapid library construction method to sequence SARS-CoV-2 gRNA

**Version 1.1**

**April 23, 2020**

#### Materials

REAGENTS

- RNaseZap (Ambion, Cat. No. AM9780)
- DNA-OFF (Takara Bio, Cat. No. 9036)
- QIAamp Viral RNA Mini Kit (Qiagen, Cat. No. 52906)
- MGIEasy rRNA removal kit (BGI, Cat. No. 1000005953)
- DNase I (RNase-free) (NEB, Cat.No.M0303)
- Tris-HCl (1M, pH 7.6; ROCKLAND, Cat. No. MB-003)
- $MgCl_2$ (1 M; Invitrogen, Cat. No. AM9530G)
- N,N-Dimethylformamide (for molecular biology, ≥99%; Sigma, Cat. No. D4551)
- DPEC-treated water (Invitrogen, Cat. No. AM9915G)
- Recombinant RNase Inhibitor (40 U/μl; Takara, Cat. No. 2313)
- Deoxynucleotide (dNTP) Solution Set (NEB, Cat. No. N0446S)
- Superscript II reverse transcriptase (Invitrogen, Cat. No. 18064014)
- DTT (0.1M; Invitrogen, Cat. No. 18064014)
- Betaine solution (5 M; Sigma, Cat. No. B0300)
- TruePrep DNA Library Prep Kit V2 for Illumina (Vazyme, Cat. No. TD501)
- PEG8000 (VWR Life Science, Cat.No.97061)
- ATP (10 mM; NEB, Cat. No. P0756)
- Q5 High-Fidelity 2x Master Mix (NEB, Cat. No. M0492)
- ChamQ SYBR qPCR master mix (Vazyme, Cat. No. Q311-02)
- VAHTS DNA Clean Beads (Vazyme, Cat. No. N411)
- Ethanol (200 proof, for molecular biology; Sigma, Cat. No. E7023)
- TargetSeq One Cov Kit (iGeneTech, Cat. No. 502002-V1)
- xGen Universal Blockers (IDT, Cat.No. 1079586)
- All oligos were acquired from Sangon.

SUPPLIES

- Millex-GP Syringe Filter Unit (0.22 μm, polyethersulfone; Millipore, Cat. No. SLGP033RB)
- 0.2 mL Thin Wall PCR Tubes (Axygen, Cat. No. PCR-02-C)

EQUIPMENT

- Thermo cycler
- Magnetic stand
- Vortexer
- Real-Time PCR machine
- A compatible Illumina DNA sequencing instrument

REAGENT SETUP

- **SARS-Cov-2 infected samples**

SARS-Cov-2 infected samples, including pharyngeal swabs, sputum samples or stool samples are collected following clinical guidelines. All of the samples, or their viral transfer media, must be deactivated at 56 °C for 30 min before nucleic acid extraction.

**IMPORTANT NOTE:** All the procedure should be operated in a BSL-3 laboratory before the total RNA is ready.

- **Total RNA**

  Total RNA can be extracted from SARS-Cov-2 infected samples by QIAamp Viral RNA Mini Kit, while omitting the addition of carrier RNA if possible. Then use MGIEasy rRNA removal kit to remove the ribosomal RNA, and DNase I to remove the DNA. The final elution volume is 12-20 μl for each sample.

  **IMPORTANT NOTE:** All the procedure should be operated in a BSL-3 laboratory before the total RNA is ready.

- **Oligo dT primer**

  Oligo dT primer (5'-$T_{30}$VN-3') anneals to all the RNAs containing a poly(A) tail. 'N' is any base and 'V' is either A, C or G. Dissolve the oligonucleotide in DPEC-treated water, to a final concentration of 100 μM. The oligo solution can be stored at −20 °C for at least 6 months.

- **Random decamer primer**

  Random decamer primer (5'-$N_{10}$-3') anneals to the RNA randomly. 'N' represents any base. Dissolve the oligonucleotide in DPEC-treated water, to a final concentration of 100 μM. The oligo solution can be stored at −20 °C for at least 6 months.

- **dNTP mix**

  Combine equal volume of dATP, dTTP, dCTP, dGTP in Deoxynucleotide (dNTP) Solution Set.

- **5X TD Buffer**

  50 mM Tris-HCl, 25 mM $MgCl_2$, 50% N,N-Dimethylformamide (DMF). This buffer can be stored at 4℃ for at least 6 months.

- **N-ch-F primer**

  N-ch-F primer (5'-GGGGAACTTCTCCTGCTAGAAT-3') is the forward primer used to amplify the N region of SARS-Cov-2 in qPCR. Dissolve the oligonucleotide in DPEC-treated water, to a final concentration of 100 μM. The oligo solution can be stored at −20 °C for at least 6 months.

- **N-ch-R primer**

  N-ch-R primer (5'-CAGACATTTTGCTCTCAAGCTG-3') is the reverse primer that used to amplify the N region of SARS-Cov-2 in qPCR. Dissolve the oligonucleotide in DPEC-treated water, to a final concentration of 100 μM. The oligo solution can be stored at −20 °C for at least 6 months.

#### Procedure

**1)** Clean the work space, including the hood and pipettes, with DNA-off and RNase-Zap. Filter the self-made buffer with 0.22 μm filter. Use a thermal cycler with a heated lid set to 105 °C for all incubations throughout this protocol. All reaction mixes should be set up on ice.

**IMPORTANT NOTE:** All the procedure should be operated in a BSL-3 laboratory before the total RNA is ready.

**Reverse transcription**

**2)** Prepare preRT mix on ice by adding 0.2 µl RNase Inhibitor (40U/µl), 0.2 µl Oligo dT (100 µM), 2 µl $(N)_{10}$ random primer (100 µM), 0.8 µl dNTP mix (25 mM) to 5.4 µl total RNA.
**NOTE:** If the volume of the RNA sample is more or less than 5.4 µl, please scale up or down the whole reaction volume proportionally.

**3)** Mix the reaction gently and thoroughly without bubbles and incubate it at 72°C for 3 min, then quickly cool it on ice.

**4)** Prepare RT mix by combining the reagents in the table below.

| Component | Volume (µl) | Final concentration |
|---|---|---|
| SuperScript II reverse transcriptase (200U/µl) | 1.00 | 200 U |
| RNase Inhibitor (40U/µl) | 0.50 | 20 U |
| SuperScript II first strand buffer (5x) | 4.00 | 1x |
| DTT (0.1M) | 1.00 | 5 mM |
| Betaine (5M) | 4.00 | 1 M |
| $MgCl_2$ (1M) | 0.12 | 6 mM |
| DPEC water | 0.78 | - |
| Total Volume | 11.40 | - |

**5)** Add 11.40 µl RT mix to the samples from step 3 and gently pipette without bubbles. Incubate the reaction at 42°C for 1.5 hour for reverse transcription, and followed by 70°C 15min to inactivate SuperScript II reverse transcriptase. Then put the sample on ice.

**Tagmentation**

**6)** Dissolve PEG8000 powder to DPEC-treated water with concentration of 40% (w/w), and filter the solution by 0.22 µm filter.

**7)** Prepare the tagmentation mix by combining the reagents listed below.

| Component | Volume (µl) | Final concentration |
|---|---|---|
| V50 (TruePrep DNA Library Prep Kit V2) | 1.00 | - |
| 5xTD buffer | 8.00 | 1x |
| RNase Inhibitor (40U/µl) | 1.00 | 40 U |
| 40% PEG8000 | 3.40 | 3.4% |
| ATP (10 mM) | 4.00 | 1.00 mM |
| DPEC water | 2.60 | - |
| Total Volume | 20.00 | - |

**8)** Add the tagmentation mix to the sample from step 5. Pipette the reaction gently and thoroughly. Incubate the reaction at 55°C for 30 min, then cool it on ice.

**Amplification and Sample Pooling**

**9)** Add 0.8 µl SuperScript II reverse transcriptase and 40.8 µl Q5 High-Fidelity 2x Master Mix to the tagmentation products.

**10)** Mix the reaction well without forming bubbles. Incubate the reaction at 42°C for 15 min to fill the 9 bp gap left by Tn5 transposome, followed by 70°C 15 min to inactivate SuperScript II reverse transcriptase.

**11)** Prepare the PCR mix by combining 4 µl N6xx index primer (10 µM), 4 µl N8xx index primer (10 µM) and 8 µl Q5 High-Fidelity 2x Master Mix. The final concentration of each index primer is around 0.4 µM.

**12)** Add PCR mix to the sample from step 10 and pipette thoroughly. Perform PCR as detailed below.

| Cycle | Denature | Anneal | Extension | Hold |
|---|---|---|---|---|
| 1 | 98°C, 30 s | - | - | |
| 2-19 | 98°C, 20 s | 60°C, 20 s | 72°C, 2 min | |
| 20 | - | - | 72°C, 5 min | |
| 21 | - | - | - | 4°C |

**13)** For targeted SARS-CoV-2 deep sequencing, continue with **qPCR and Sample Pooling.** For metagenomic sequencing, pool 8-16 libraries in equal volumes and go to **Step 18**.

##### qPCR and Sample Pooling

**14)** Take 1 µl PCR product and dilute it 200-fold with nuclease-free water.

**15)** Prepare qPCR mix by adding 0.05 µl N-ch-F primer (100 µM), 0.05 µl N-ch-R primer (100 µM), 5 µl ChamQ SYBR qPCR master mix and 3.9 µl nuclease-free water to 1 µl diluted template.

**16)** The program of qPCR is performed at 95°C for 60s, followed by 40 cycles of [95°C 5s, 60°C 15s].

**17)** Pool 8-16 libraries together according to their Ct values as following:

| Ct | Volume taken (µl) |
|---|---|
| >28 | 50.0 |
| 24-28 | 16.0 |
| 20-24 | 3.0 |
| <20 | 0.5 |

##### Purification

**18)** Place VATHS DNA clean beads at room temperature for 15 min, then vortex violently.

**19)** Add equal volume of beads (1x) to the pooled samples and vortex violently. Incubate the mixture at room temperature for 5 min.

**20)** Transfer the tube to compatible magnetic stand until the solution is clear.

**21)** Carefully remove the solution without disturbing beads.

**22)** Wash beads with 200 µl 80% ethanol (freshly prepared) and incubate for 30 s, then remove the ethanol. Repeat this step one more time.

**23)** Dry the beads on magnetic stand with cap open until the color of beads gets light. Add 52 µl nuclease-free water and close the cap, vortex violently to wash DNA off.

**24)** Incubate the tube at room temperature for 5 min off the magnet stand.

**25)** Quickly spin down the tube then place it on magnetic stand until the solution is clear.

**26)** Carefully aspirate 50 µl supernatant to a clean tube without disturbing beads. The library can be restored at -20°C for 6 months.

**IMPORTANT NOTE:** The equal-volume mixed libraries are now ready for sequencing.

**SARS-Cov-2 Sequence Enrichment**

**27)** Subject the purified Ct-adjusted library pool to one round of SARS-Cov-2 sequence capture following the instruction of TargetSeq One Cov Kit. Replace the iGeneTech Blocker with the IDT xGen Universal Blockers.

**High-Throughput Sequencing**

28) Both the metagenomic library and the SARS-Cov-2 targeted library can be sequenced on any Illumina platform with paired-end mode. 10-20 million reads are typically collected.
